# Exploring RNA modifications in infectious non-coding circular RNAs

**DOI:** 10.1101/2024.03.12.584625

**Authors:** Pavel Vopalensky, Anton Škríba, Michela Chiumenti, Lucia Ďuričeková, Anna Šimonová, Ondřej Lukšan, Francesco di Serio, Beatriz Navarro, Hana Cahova

## Abstract

Viroids, small circular non-coding RNAs, act as infectious pathogens in higher plants, demonstrating high stability despite consisting solely of naked RNA. Their dependence of replication on host machinery poses the question of whether RNA modifications play a role in viroid biology. Here, we explore RNA modifications in the avocado sunblotch viroid (ASBVd) and the citrus exocortis viroid (CEVd), representative members of viroids replicating in chloroplasts and the nucleus, respectively, using LC–MS and Oxford Nanopore Technology (ONT) direct RNA sequencing. Although no modification was detected in ASBVd, CEVd contained approximately one m^6^ A per RNA molecule. ONT sequencing predicted three m^6^ A positions. Employing orthogonal SELECT method, we confirmed m^6^ A in two positions A353 and A360, which are highly conserved among CEVd variants. These postitions are located in the left terminal region of the CEVd rod-like structure where likely RNA Pol II and and TFIIIA-7ZF bind, thus suggesting potential biological role of methylation in viroid replication.

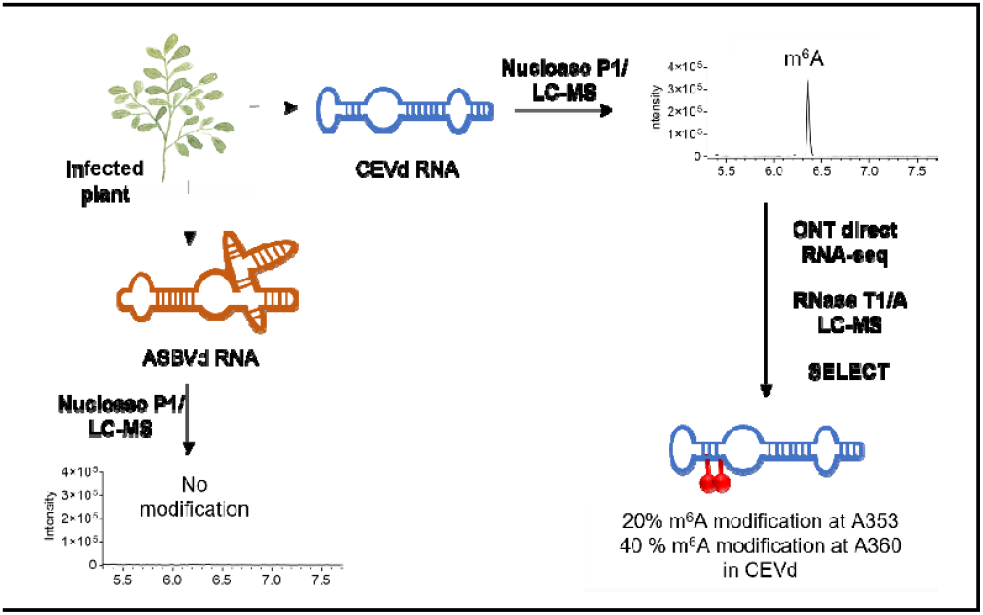

---

RNA is a key molecule in all cellular processes and plays a plethora of roles. The functional information in RNA is encoded in three layers: sequence, structure and chemical modifications. To date, there are approximately 170 RNA modifications discovered across all domains of life^1^. The role of RNA modifications is well described for abundant RNA species such as rRNA^2^ and tRNA^3, 4^; however, it is not as well understood in other RNA types (e.g. mRNA, regulatory RNA and viral RNA)^5, 6^. The low abundance of these RNAs and contamination from rRNA and tRNA hinders their in-depth analysis^7, 8^. Viruses are good model systems for studying RNA modifications thanks to their intrinsically simple organization and amplification in infected cells^9^. So far, the best-described modifications in viral RNA are 6-methyladenosine (m^6^A)^10, 11^⍰and 2⍰-O-methylnucleoside (Nm)^12^, which have been detected, for example, in HIV-1. Other modifications, such as 1-methyladenosine (m^1^A) or 5-methylcytidine (m^5^C), were detected rather in viral co-packed tRNA in HIV-1^13^⍰ and tRNA fragments in members of the *Picornavirales* ^14^.

Subviral agents infecting plants, such as viroids, offer even better opportunities to identify new RNA modifications and their potential biological roles. Viroid RNA does not code for proteins and the catalytical activity needed for their infectivity is provided either by their own ribozyme structure (family *Avsunviroidae*) or by host enzymes, mimicking structural features of host nucleic acid. The replication of viroids can take place either in the nucleus (family *Pospiviroidae*) or in chloroplasts (family *Avsunviroidae*)^15^ (Figure 1A). Despite being naked circular RNAs of small size (250–450 nt), viroids are highly stable in natural environments. The stability and intimate relationship between sequence, structure and function poses the question whether RNA modifications play a role in viroid biology.

**Figure 1.**
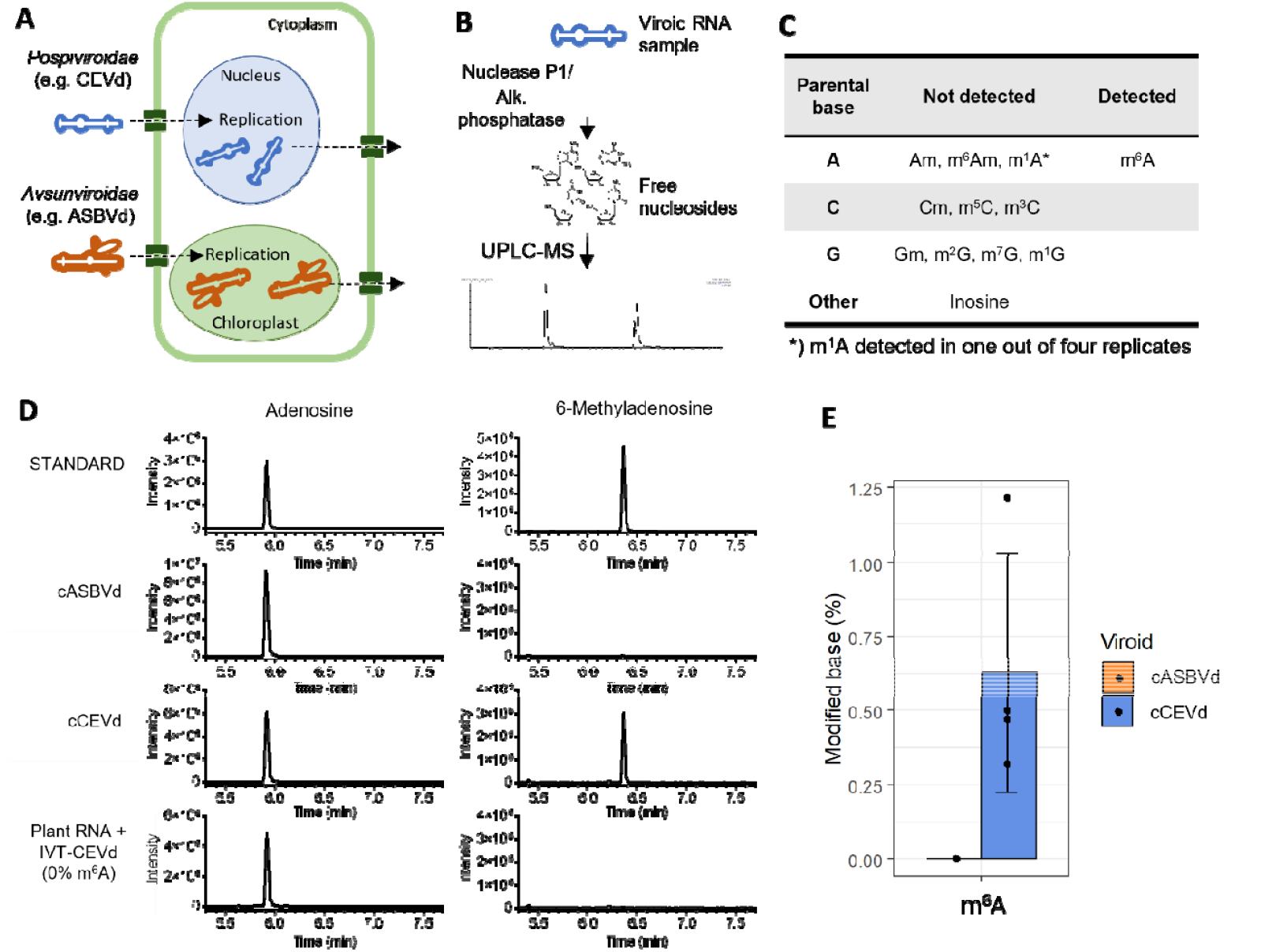
Identification of RNA modifications in viroid RNA. (A) The citrus exocortis viroid (CEVd, family Pospiviroidae) replicates in the nucleus whereas the avocado sunblotch viroid (ASBVd, family Avsunviroidae) replicates in chloroplasts. (B) Double PAGE-purified viroid circular RNA sample (cCEVd, cASBVd) was digested by nuclease P1 and alkaline phosphatase into nucleoside form. The resulting mixture was subjected to LC–MS analysis. (C) Out of eleven types of RNA modification screened for, only m^6^A was reproducibly detected in cCEVd RNA. (D) LC–MS extracted ion chromatograms of adenosine and 6-methyladenosine from nuclease P1-digested cASBVd, cCEVd and IVT-CEVd (negative control) RNA spiked in the RNA of healthy plants. (E) Quantification of m^6^A in cASBVd and cCEVd RNA samples (measured in biological triplicates). The error bar represents the standard deviation; the black dots are individual values from each replicate. The y-axis represents the percentage of m^6^A across all adenosines within the RNA sequence.

So far, attempts to identify RNA modifications in viroids have employed the bisulfite sequencing, which did not reveal any m^5^C in viroid RNA^16, 17^ ⍰. Additionally, bioinformatic prediction was performed, but without further experimental confirmation^18^. To date, however, no systematic search for RNA modifications in viroids has been reported. In this study, we investigated the presence of the most common RNA modifications^7^ in two representative members of both viroid families by employing liquid chromatography–mass spectrometry (LC–MS) analysis, Oxford Nanopore Technology (ONT) direct RNA sequencing and an elongation- and ligation-based qPCR amplification method (SELECT). Whereas the avocado sunblotch viroid (ASBVd, *Avsunviroidae*) did not contain any of the RNA modifications screened for, we did detect the m^6^A methylation in the citrus exocortis viroid (CEVd, *Pospiviroidae*).

## Results and Discussion

Viroids are classified into the families *Avsunviroidae* and *Pospiviroidae*, which include members replicating in the chloroplasts and in the nucleus, respectively (Figure 1A). We selected the viroids ASBVd (*Avsunviroidae*) and CEVd (*Pospiviroidae*), which are known to accumulate at high levels in their natural or experimental host plants (*Persea americana* and Gynura aurantiaca, respectively) as representative members of each family to screen for modifications in viroid RNAs. We carried out our investigation on viroid circular RNA forms isolated from tissues infected with these viroids by applying a purification protocol consisting of the following two steps: (i) preparation of extracts enriched in highly structured RNAs by chromatography on CF11 cellulose in the presence of 35% EtOH^19^ and (ii) separation of CF11 RNAs by double polyacrylamide gel electrophoresis (PAGE) consisting of a non-denaturing electrophoresis followed by a denaturing electrophoresis. The latter electrophoresis separated linear RNAs from circular viroid RNAs that could be easily recovered from the gel. This approach allowed us to obtain highly purified viroid circular RNA, reducing contamination by host RNA (Figure S1). After digestion of the gel-purified circular viroid RNAs (hereinafter referred to as cCEVd and cASBVd) by nuclease P1 and alkaline phosphatase into nucleosides, LC–MS analysis (Figure 1B) was performed to screen for the eleven most common modifications, namely 2⍰-O-methyladenosine (Am), m^6^A, 6-methyl-2⍰-O-methyladenosine (m^6^Am), m^1^A, 2⍰-O-methylcytidine (Cm), m^5^C, 3-methylcytidine (m^3^C), 2⍰-O-methylguanosine (Gm), 2-methylguanosine (m^2^G), 7-methylguanosine (m^7^G) and inosine (I) (Figure 1C, Figure S2, Table S6). Out of these, we observed m^6^A RNA modification signal in cCEVd-digested RNA (Figure 1D–E). This signal was observed in all four replicates tested. By contrast, no RNA modification was detected in the digested cASBVd.

To exclude the possibility that m^6^A comes from a host RNA molecule co-migrating with CEVd RNA, we performed a parallel control experiment. We prepared *in vitro*-transcribed CEVd RNA (IVT-CEVd) using *in vitro* transcription by T7 RNA polymerase (see the supplementary material) with diadenosine diphosphate (Ap_2_A) included in the mixture to serve as a 5⍰ RNA cap. Uncapped RNA was degraded using 5⍰ polyphosphatase and Terminator exonuclease. The presence of the Ap_2_A RNA cap enables efficient ligation (circularization) by truncated T4 RNA ligase 2. The non-circularized forms remaining after ligation were degraded using RNAse R, resulting in pure circular viroid RNA. IVT-CEVd migrates exactly as cCEVd from infected tissue but does not contain any RNA modification. Therefore, IVT-CEVd was spiked in the RNA (the fraction consisting of highly structured RNAs) isolated from non-infected *Gynura* plants and subsequently recovered by the same double PAGE procedure used to isolate cCEVd from infected plants (Figure S1). LC–MS analysis of this IVT-CEVd RNA did not detect any m^6^A, thus excluding the possibility that the m^6^A signal observed in cCEVd RNA came from co-migrating host RNA (Figure 1D).

The quantified amount of detected m^6^A corresponds to 0.5–1.2% of all adenosines (75 A in the 371-nt-long CEVd RNA), which would approximately mean 0.5–1 m^6^A per one molecule of CEVd RNA. Given this fact, we decided to explore whether this modification was present in a specific position or whether it was distributed over the viroid RNA sequence. Because traditional sequencing methods used for the identification of m^6^A position do not provide single-nucleotide resolution or require a large amount of input material (e.g. MeRIP^20^, m^6^A-seq^21^ or miCLIP protocol^22^), we employed ONT direct RNA sequencing in combination with published m^6^A prediction algorithms to gain single-nucleotide resolution^23^. A similar method was used recently for studies of m^6^A in two types of viruses^24^. We used IVT-CEVd to establish the ONT sequencing protocol and to provide a control dataset required by most prediction algorithms (Figure 2A). First, we optimized the ratio of Zn^2+^ ions needed to RNA for a single cleavage site to gain mostly full-length reads (Figure 2B)^25^. The linearized RNA was polyadenylated and sequenced employing the ONT RNA002 and 9.4.1 flow cell or ONT RNA004 and RNA flow cell, producing at least 29 thousand CEVd-mapping reads (Figure 2C). The three IVT-CEVd samples were sequenced using two different templates starting from two different positions within the viroid sequence (Figure S3, see the supplementary material).

**Figure 2.**
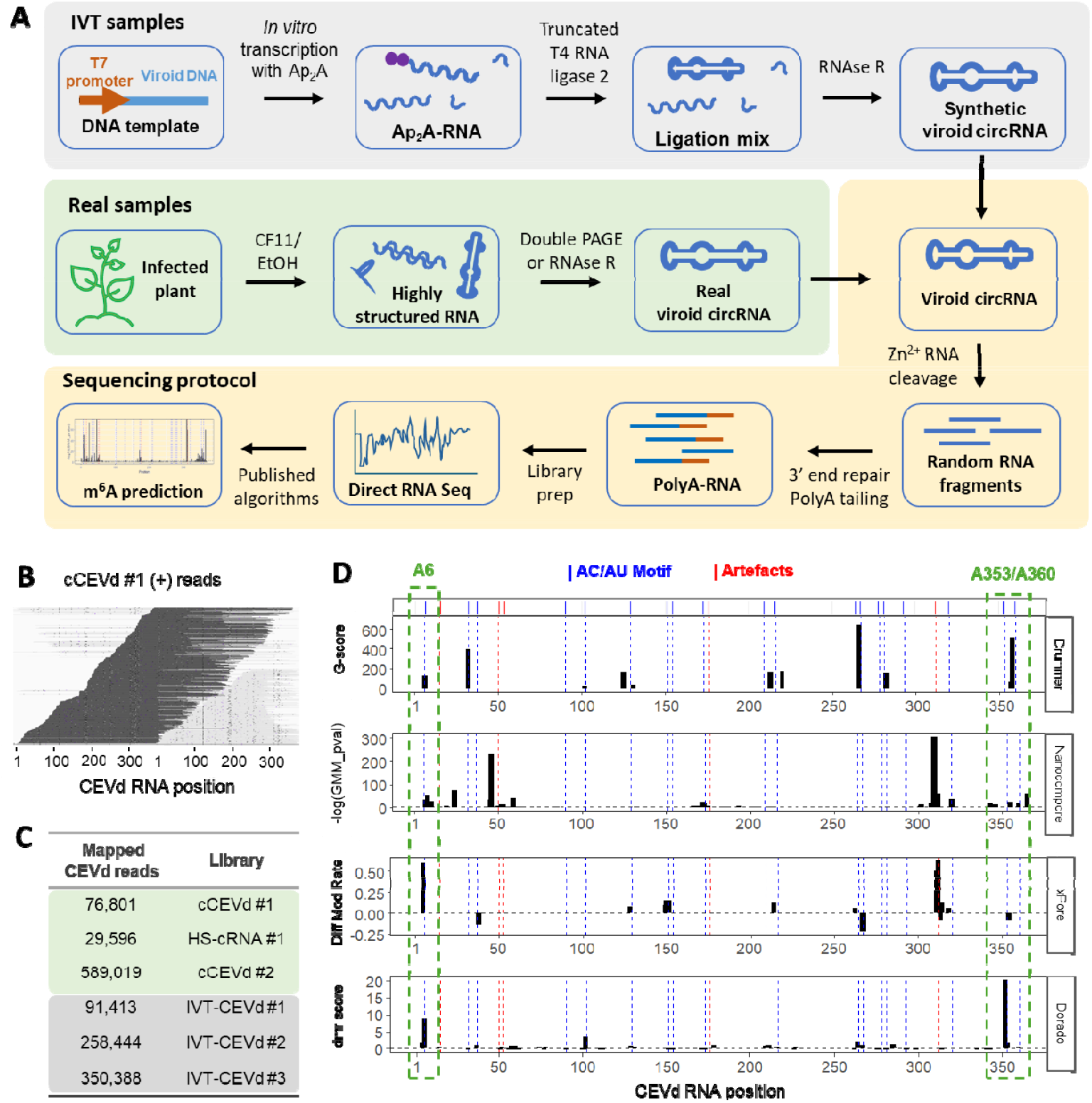
Identification of m^6^A position by ONT direct RNA sequencing. (A) Overview of the sample preparation procedure (IVT-CEVd samples – grey background, CEVd samples from plant tissues – green background, sequencing protocol – yellow background.) (B) Alignment of CEVd reads from a typical run against the reference CEVd sequence, visualized using Integrative Genomics Viewer^24^; the reference sequence is dimerized to enable the visualization of reads mapped to a circular sequence. A uniform read length of ∼_370 nt was observed, corresponding to CEVd RNA linearized by a single cleavage. Distributed read starts_/_ends along the reference sequence suggests a random position of circular RNA cleavage. (C) ONT sequencing reads mapping to the CEVd sequence for each experiment (see also Table S7). (D) Prediction of m^6^A sites by multiple independent algorithms. The blue dashed lines indicate the sites of the AC_/_AU dinucleotide. The red dashed lines indicate the positions of SNPs and “scars” causing high falsely positive m^6^A prediction scores at several positions (see Figure S4 for details). DRUMMER, Nanocompore and xPore prediction was based on cCEVd #1, HS-cRNA #1 / IVT-CEVd #1 and #2 samples sequenced with RNA002 chemistry whereas Dorado m^6^A prediction was performed with cCEVd #2 / IVT-CEVd #3 samples sequenced with RNA004 chemistry. The candidate sites A6, A353 and A360 (indicated by the green dashed boxes) were selected based on their scoring agreement across all four prediction approaches.

Sequencing of samples from CEVd-infected plants was performed using gel-purified cCEVd, previously analysed by LC–MS (cCEVd #1) and a highly structured fraction of RNA obtained by CF-11 chromatography and treated with RNase R (HS-cRNA #1). Importantly, cCEVd is expected to contain highly purified circular viroid RNAs, HS-cRNA is only enriched in circular RNAs and contains also host RNAs. Both samples were sequenced using the same procedure.

To predict the position of m^6^A in CEVd RNA, we applied several published algorithms (DRUMMER^26^ v1.0, xPORE^27^ v2.1 and Nanocompore^28^ v1.0.4) on four datasets (cCEVd #1, HS-cRNA #1 and the negative controls IVT-CEVd #1 and #2). For each algorithm, we set parameter thresholds based on a recent benchmarking study^23, 29^. We observed single-nucleotide polymorphisms (SNPs) due to the naturally occurring variability of viroid populations^30^ and several artefacts, such as “circularization scars”, caused by different non-templated nucleotides at the 3⍰ end of *in vitro* transcripts (Figure S4). After filtering out these artefacts, we identified three potential sites of m^6^A, namely positions A6, A353 and A360 (Figure 2D). We also used the recently released ONT RNA004 technology to sequence other double PAGE-purified circular CEVd RNA sample from infected plants (cCEVd #2) and a control IVT-CEVd (IVT-CEVd #3) (Table S7). The analysis of this ONT sequencing with Dorado (v1.0; https://github.com/nanoporetech/dorado/), predicted two of these three positions as the most frequently modified ones (Figure 2D). Interestingly, only 0.3% of all reads belonged to host plant RNAs in the two double PAGE-purified CEVd samples, cCEVd #1 and cCEVd #2, confirming the purity of the samples and providing further evidence that m^6^A must come from cCEVd RNA (Table S7).

To confirm the predicted positions of m^6^A, we employed sequence specific cleavage by specific RNases combined with LC–MS of the resulting oligonucleotides. The cCEVd was digested by RNase T1 or RNase A, which cleave after a G and a U/C nucleotide, respectively (Figure 3A). This digestion generates a mixture of oligonucleotides, the composition of which is easy to predict from the CEVd sequence variant (GenBank ID: PP446493, infecting the source plants of *G. aurantiaca*) (Figure 3B, Table S8-S9). The sensitivity of LC–MS decreases with the length of the oligonucleotide. In fact, no signal of oligonucleotides (methylated or non-methylated) longer than 8 nt was detectable in an IVT-CEVd transcript prepared with 70% of m^6^ATP/30% ATP in IVT mixture (IVT-CEVd 70% m^6^A) (Figure 3C–D, Figure S5). However, digestion with at least one of the two RNases produces at least one oligonucleotide with a maximum length 7 nt from each AU or AC motif, which is the core of the recognition sequence of m^6^A RNA methyltransferases in plants^31^ (Figure 3B), allowing to ultimately confirm or exclude the presence of the m^6^A methylation at each position of interest. First, we focused on predicted positions (A6, A353 and A360) and then we searched for the mass of respective oligonucleotides in their unmethylated and monomethylated versions. As standards we used IVT-CEVd 70% m^6^A digested in the same way as the gel-purified CEVd from infected tissues (cCEVd) (Figure 3C-D, Figure S5). Surprisingly, we did not observe any signal of methylated oligonucleotides containing any of the three predicted positions in the cCEVd RNA sample (Figure 3C-D, Figure S5). We therefore extended the analysis to all oligonucleotides containing AU or AC motif (Figure 3B, Figure S6)^31^. Also, in this case, we did not identify any methylated oligonucleotides in our analysis (Figure S5). Extending the search for methylation to all other adenosine-containing oligonucleotides (data not shown) did not reveal the presence of m^6^A either. This finding suggests that the m^6^A detected by LC–MS, using nuclease P1-digested cCEVd RNA, is not present at a single position in the viroid but might instead be distributed at lower stoichiometries at multiple specific positions (presumably those predicted by direct ONT sequencing). Considering the sensitivity of our RNase/LC–MS method, we can estimate the limit of detection from the signal-to-noise ratio (S/N = 3). For example, at position 360, methylation would be detected if more than 30% of the AGAC sequence contains m^6^A modification. It is also important to mention that the limit of detection is better with shorter sequences and vice versa.

**Figure 3.**
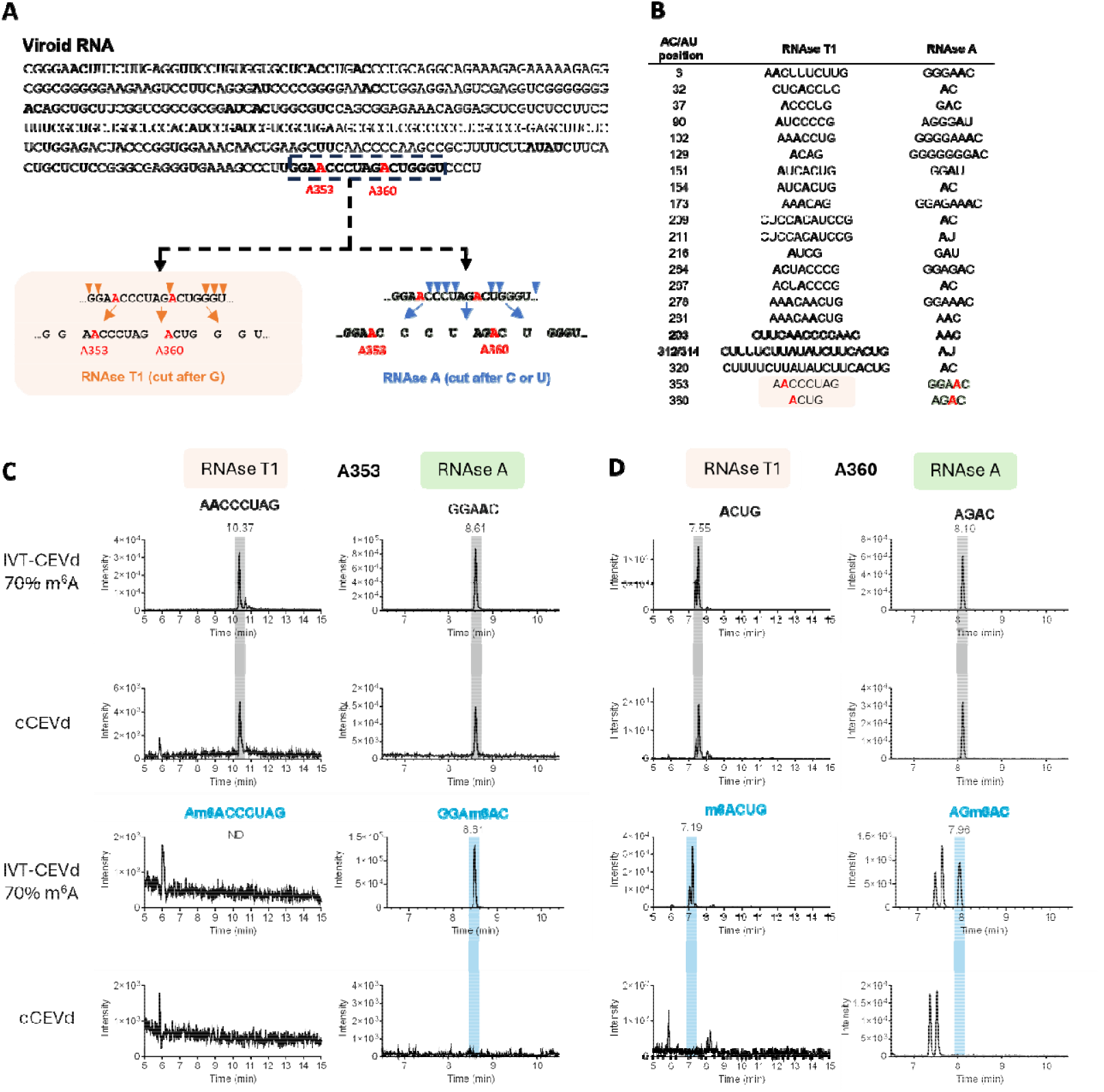
Verification of candidate m^6^A positions by the RNAs ⍰ /⍰ LC–MS approach. (A) Purified circular form of CEVd (cCEVd or IVT-CEVd) is digested by RNase T1 or RNase A, which cleave the RNA after each G or C/U nucleotide, respectively, generating a mixture of specific short oligonucleotides for LC–MS analysis. Given the different specificity of the two enzymes, each candidate position is contained in a unique combination of oligonucleotides from the two digests. (B) The sequences of the resulting RNA oligonucleotides from digesting the CEVd RNA. Only oligonucleotides containing the AC/AU dinucleotide motif are listed. Note that the specific site of the modified A position could be identified by the combination of both RNases. However, for example in the cases of A264 and A267, which are contained in the same oligonucleotide produced by RNase T1, they could be distinguished by two different oligonucleotides produced by RNase A digestion. (C) Representative LC–MS extracted ion chromatograms of unmodified (peaks highlighted by grey box) and m^6^A-modified (peaks highlighted by blue box) oligonucleotides at candidate sites A353 and A360. IVT CEVd (0 and 70% m^6^A) was used as a positive control. For extracted ion chromatograms of all other positions listed in Fig. 3B, see Table S8-S9 Figure S5.

Because the sensitivity of the RNase/LC-MS method did not allow the detection of the m^6^A at the candidate sites, we turned to an orthogonal method - single-base elongation- and ligation-based qPCR amplification method (SELECT) ^32^. In this method, two primers are annealed to RNA template adjacent to m^6^A site with one nucleotide gap. The gap is filled by Bst 2.0 DNA polymerase and the elongated primer is subsequently ligated to the second one by splint ligation employing SplintR ligase. Both enzymes are less efficient in the presence of m^6^A in the RNA template (Figure 4A). This method is suitable for detecting m^6^A at candidate positions if the surrounding ∼20 nucleotides are known and if RNA samples with various percentage of m^6^A at the given candidate position are available to construct the calibration curve. We synthetized IVT cCEVd containing 0% and 100% m^6^A and mixed these in various proportions to obtain samples with 100, 50, 25, 12.5 and 0% m^6^A. The exact percentage of m^6^A in IVT cCEVd RNA was determined by LC-MS measurement (Figure S7 A). Using these calibration samples, we determined that, in the double-PAGE purified cCEVd, ∼ 20% of the A353 position and ∼ 40% of the A360 position contain m^6^A. However, we did not detect any m^6^A at the candidate position A6 (Figure 4B and Figure S7 B-C). The inability to detect m^6^A at A6 position might be due to the downstream low-complexity sequence context of the A6, with ∼ 47% and ∼ 71% GC content within the upstream and sequences (see Figure 3A), respectively, rendering the SELECT method unsuitable for this region. This fact might be reflected by the poor performance of the calibration curve for A6. On the other hand, the analysis of 148 CEVd sequences from public databases revealed that the adenosine at position 6 is present in less than 10% of them, with the vast majority possessing a U at this position (>80%) (Figure S6 B), supporting the observation that the A6 position is probably not functionally important and, when present, is not methylated. In contrast, A353 and A360 are highly conserved positions in all the CEVd variants, a finding consistent with a potential biological role of the observed methylated nucleotides at these positions. In addition, we explored the conservation of the A353 and A360 among the other species of the genus *Pospiviroid* to which CEVd belongs and we observed that motif GGNACC (N = A or AA or deletion), surrounding CEVd A353, is conserved in the terminal left region of the rod-like conformation of all these viroids (Figure 4C and Figure S8). In this region of the potato spindle tuber viroid (PSTVd), the type species of the genus, the binding domain of RNA Pol II and TFIIIA-7ZF, two proteins involved in replication of nuclear viroids, was also mapped^33, 34^. Regarding the position A360, while in the CEVd variant studied here it is a part of an AC motif (A360-C361), other reported variants showed polymorphisms in position 361 (70% U, ∼28% G, and 2% C).

**Figure 4:**
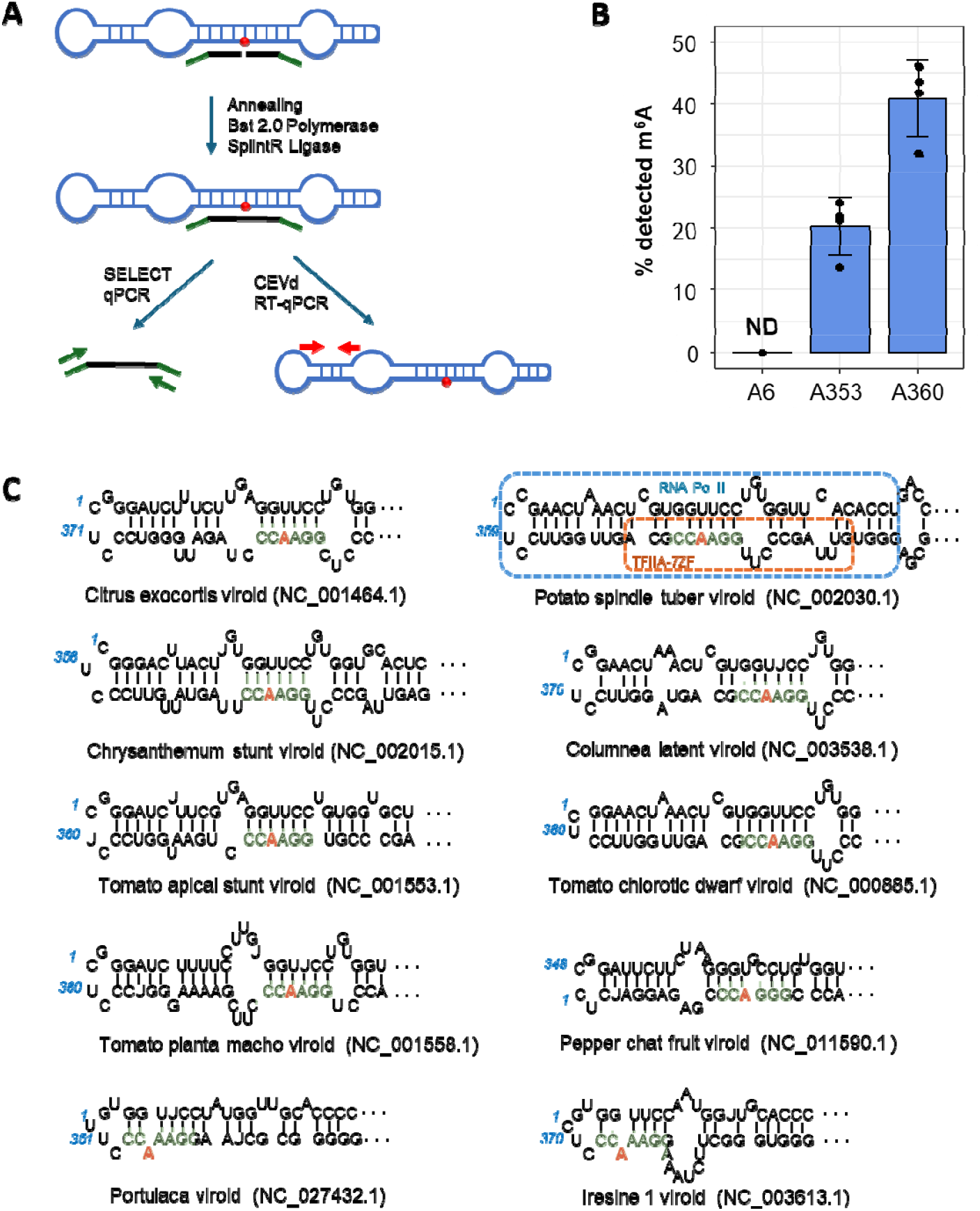
Detection of m^6^A at candidate positions within the CEVd sequence by SELECT and their conservation within pospiviroids. (A) The principle of the SELECT method: Two anti-sense oligonucleotides spanning the candidate m^6^A are annealed to the viroid RNA. Subsequently, the Bst2.0 polymerase is used to fill-in the thymidine nucleotide complementary to the candidate m^6^A site and the SplintR Ligase ligates the adjacent oligonucleotides into one fragment of ssDNA (SELECT product). As both Bst 2.0 polymerase and SplintR ligase enzymatic activity is lower in the presence of m^6^A compared to A, the resulting amount of ligated ssDNA fragment is inversely proportional to the percentage of m^6^A at given candidate position. The amount of ligated SELECT product is estimated by qPCR and normalized to the total amount of viroid RNA using RT-qPCR with CEVd-specific set of oligos. (B) The amount of methylated adenosine at candidate sites A6, A353 and A360. Whereas we were not able to detect m^6^A on the A6 position, the position A353 and A360 showed 20% and ∼40% methylated adenosine, respectively. The black dots represent individual measured values; the error bar represents the standard deviation of the mean. (C) Primary and secondary structure of the terminal left region of the rod-like conformation of the members of the genus *Pospiviroid* (the refseq variant of each species and the ID number are in indicated below the structure). The conserved motif GGNACC has a green background, and the conserved A (A353 in citrus exocortis viroid) is highlighted red. In the secondary structure of potato spindle tuber viroid, the binding regions of RNA pol II and TFIIA-7ZF are marked by a blue and a red square, respectively.

## Conclusions

In summary, we explored the presence of RNA modifications in two viroids that are representative members of the two viroid families, employing LC–MS, ONT RNA sequencing and SELECT methods. Although we did not detect any of the most common RNA modifications in chloroplast-replicating ASBVd, we observed m^6^A in the nucleus-replicating CEVd.

This finding is in agreement with the fact that m^6^A RNA monomethyltransferases are mainly reported to occur in the nucleus^35^ but have not been described in chloroplasts⍰^36^. LC–MS analysis revealed 0.5–1 m^6^A per CEVd RNA molecule, indicating that m^6^A might be present at one specific position. To identify the position of m^6^A, we employed ONT direct sequencing in combination with several m^6^A prediction algorithms. These algorithms identified three candidate positions. We tried to confirm the methylation at these positions employing RNase-specific cleavage followed by LC–MS analysis but the sensitivity of the method was not sufficient to provide conclusive results. Therefore, we used an alternative method called SELECT confirming the methylation of two out of the three predicted candidates. The modification rate 20% m^6^A at A353 and 40 % m^6^A at A360 is in agreement with the results of the initial LC-MS analysis showing 0.5–1 m^6^A per CEVd RNA molecule. The distribution of the rather low percent of methylation among the two sites also explains why the RNase T1-⍰/⍰RNase A - based method failed to detect these modifications.

Interestingly, the A353 and A360 are positioned within the left-terminal region of the CEVd rod-like conformation. In another nuclear viroid, PSTVd, this region has been identified as the binding site of the RNA Pol II ^34^ and TFIIIA-7ZF ^33^ involved in viroid replication^15^. Further studies might elucidate the functional role of the m^6^A modification at these positions for the entire viroid infectious cycle.

## Author Contributions

P.V., A.Š., B.N., F.d.S. and H.C. designed the experiments and coordinated the project. P.V., A.Š., L.D., A.Š., and H.C. performed the experiments. P.V., M.C. and O.L. performed all bioinformatic analyses. H.C. a B.N. supervised the work. P.V., F.d.S., B.N. and H.C. wrote the paper.

## Acknowledgement

This work was funded as a bilateral scientific project between the Italian Research National Council (CNR) and the Czech Academy of Sciences (CAS) under the joint programme for the 2022–2024 triennium (CNR-22-21). We also acknowledge funding from the Operational Programme Johannes Amos Comenius (OP JAC) project RNA for Therapy, reg. No. CZ.02.01.01/00/22_008/0004575 co-financed by the EU and the European Research Council Executive Agency (ERCEA) under the European Union’s Horizon Europe Framework Programme for Research and Innovation (grant agreement No. 101041374 – StressRNaction). Views and opinions expressed are however those of the authors only and do not necessarily reflect those of the European Union. Neither the European Union nor the granting authority can be held responsible for them.

## Data Availability

The sequencing data that support the findings of this study are openly available in ENA-European Nucleotide Archive at https://www.ebi.ac.uk/ena/browser/view/PRJEB75479, project accession number: PRJEB75479 and Zenodo under the DOI: 10.5281/zenodo.14418523

