## Supplementary Data for "Exploring RNA modifications in infectious non-coding circular RNAs"

**Supplementary materials and methods**

**Plant material and RNA isolation**

Leaves of healthy and CEVd-infected *Gynura aurantiaca* plants and of an ASBVd-infected avocado (*Persea americana* Mill.) tree were used as source for RNA isolation. Nucleic acid preparations enriched in highly structured RNAs were obtained by extracting leaves (20 g) with buffer-saturated phenol and partitioning the nucleic acids by chromatography on non-ionic cellulose CF-11 (Whatman) with 1x STE [50 mM Tris-HCl, pH 7.2, 100 mM NaCl, 1 mM ethylenediaminetetraacetic acid (EDTA)] containing 35% ethanol as described by Pallás et al., 1987^1^. The RNA preparations enriched in highly structured RNAs (HS-RNA) were recovered by ethanol precipitation and resuspended in H_2_O (1 ml).

**Purification of viroid circular RNAs**

Purification of circular RNA forms of viroids was performed by fractionation of the RNA preparations enriched in highly structured RNAs in two sequential 5% polyacrylamide gel electrophoresis (PAGE) under non-denaturing conditions followed by denaturing conditions as described in Flores et al., 1985^2^ (Figure S1). Briefly, the HS-RNA (30 µL per well) were first loaded in a non-denaturing 5% PAGE in 1xTAE (40 mM Tris, 20 mM sodium acetate and 1 mM EDTA, pH 7.2 with acetic acid) buffer. After staining with ethidium bromide, about a 0.8 cm segment of the gel containing the band migrating according to the viroid size (citrus exocortis viroid, CEVd: 370 nt; avocado sunblotch viroid, ASBVd: 246 nt) was cut and applied directly on top of the denaturing 5% PAGE in buffer 0.25X TBE [22.5 mM Tris, 22.5 mM boric acid and 0.5 mM EDTA (pH 8·3)], containing 8 M urea in the gel^3, 4^. Under denaturing conditions, circular RNAs migrate more slowly than their corresponding linear forms. Therefore, after denaturing PAGE and ethidium bromide staining, the bands corresponding to the circular forms of the viroid migrate in the upper region of the gel (Figure S1). The portion of gel containing such a band was cut and the RNA eluted from the gel by grinding with a mixture (1:1) of water-saturated phenol and buffer (100 mM Tris-HCl pH 8.9, 1 mM EDTA, 0.5 % SDS)^1^. The nucleic acids were recovered from the aqueous phase by ethanol precipitation.

**Cloning of full-length CEVd viroid cDNA**

To generate the constructs pGemTeasy-CEVd-IVT#1 and pGemTeasy-CEVd-IVT#2 (Figure S3), used as template for the synthesis of *in vitro* monomeric transcripts (IVT) of CEVd, monomeric cDNAs of the CEVd variant infecting *G. aurantiaca* (GenBank ID PP446493) containing the T7 RNA polymerase promoter sequence at 5’ end, were cloned in pGemT-easy vector (Promega). To this aim, cDNA synthesis was performed with reverse transcriptase Superscript IV (Invitrogen) in 20 µl of reaction volume using 9 µl of CF11-RNA preparation from CEVd-infected *G. aurantiaca* plants and random hexamers, following the supplier conditions. 2µl of the cDNA reaction were used for PCR amplification using two different CEVd specific primer pairs (CEVd2.5phi-For1/CEVd2.5phi-Rev2 and CEVd2.5phi-For3/ CEVd2.5phi-For4, Table S1) able to amplify full-length monomeric cDNAs of CEVd, with the 5’ end located in two different positions of the CEVd genome (Table S1). The CEVd forward primers contained at the 5’ end an overhang sequence coding for the T7 class II promoter phi2.5 (TAATACGACTCACTATT). The PCR amplicons were cloned in pGemTeasy vector and their sequence was confirmed by sanger sequencing.

**In vitro transcription (IVT) of viroid RNA**

The template DNA was prepared by PCR amplification from plasmids pGemTeasy-CEVd-IVT#1 and pGemTeasy-CEVd- IVT#2 (Figure S3), using Phusion HF (New England Biolabs #M0530). The T7 promoter was introduced to the template by using forward primer with the T7 promoter sequence. The remaining plasmid DNA was removed by digesting with *DpnI* (NEB #R0176) for 30 minutes at 37 °C. The PCR product was purified using Monarch PCR and DNA Cleanup Kit (NEB # T1030). The in vitro transcription mix (typically 50 µL) contained 20 ng/μL of template DNA, 1× reaction buffer for T7 RNAP and 62.5 units of T7 RNA polymerase (NEB, #M0251), 1 mM UTP, 1 mM CTP, 1 mM GTP, and 1 mM ATP [or 0.3 mM ATP and 0.7 mM m^6^ATP for obtaining IVT-CEVd with 70% of m^6^A (IVT-CEVd 70% m^6^A] and 1.6 mM diadenosine diphosphate (Ap_2_A) (Jena Bioscience #NU-936-5). The buffer was further supplemented by 5% dimethyl sulfoxide (DMSO), 0.12% Triton X-100, 12 mM dithiothreitol (DTT) and 4.8 mM MgCl_2_. The transcription reaction was incubated for 2 h at 37 °C. Afterwards, DNase I was added to digest the DNA template. For a total of 50 μL transcription mixture, 6 μL of 10× DNase I reaction buffer and 8 units of DNase I (NEB #M0303) were incubated at 37 °C for 60 min. Afterwards, EDTA was added to a final concentration of 50 mM and the enzymes were heat deactivated at 75 °C for 10 min followed by immediate cooling on ice. The transcription mixture was purified using RNA Clean and Concentrator™ 5 (Zymo Research, #R1016) and eluted in 25 μL water. To remove the RNA that did not carry the Ap_2_A 5’ cap, the RNA was subsequently incubated with RNA 5′-polyphosphatase and Terminator™ 5′-phosphate-dependent exonuclease. First, the purified IVT RNA was treated with 20 units of 5′-polyphosphatase (Lucigen #RP8092H) in the solution of 1× buffer in a total volume of 30 μL for 1 h at 37 °C to convert triphosphate 5’ RNA end to monophosphate. Afterwards, the RNA was purified by RNA Clean and Concentrator™ 5 (Zymo Research, #R1016) and eluted in 25 μL of water. 2 units of Terminator™ 5′-phosphate-dependent exonuclease (Lucigen #TER51020) and 3.3 μL of buffer were then added and the mixture was incubated at 30 °C for 1 h to degrade monophosphate 5’end-carrying RNA. The resulting RNA carrying only 5’ Ap_2_A cap was purified using RNA Clean and Concentrator™ 5 (Zymo Research).

**Circularization of in vitro transcribed viroid RNA**

To circularize the Ap_2_A RNA, the RNA was ligated using T4 RNA Ligase 2, truncated (NEB #M0242) that requires the pre-adenylated (i.e. Ap_2_A) 5’ end of RNA for ligation to occur (Figure S3). The reaction mixture containing 1x T4 RNA Ligase Reaction Buffer, 10% PEG8000, 3 μg Ap_2_A-RNA and 10 U/μL truncated T4 RNA Ligase 2 was incubated 16 h at 16 °C. Afterwards, the RNA was purified using RNA Clean and Concentrator™ 5 (Zymo Research) and non-circular RNA was removed by incubating with 2 units of RNase R (Lucigen #RNR07250) in supplied buffer for 2 hours at 37 °C. The resulting circular RNA was purified by RNA Clean and Concentrator™ 5 (Zymo Research).

**Nuclease P1 digestion**

Purified circular forms of ASBVd (cASBVd), CEVd (cCEVd), and IVT-CEVd (200-300 ng each) RNAs were digested to nucleotides with 100 units of Nuclease P_1_ enzyme (NuP1, New England BioLabs, #M0660S) in 50 mM ammonium acetate (Sigma-aldrich, #A1542) buffer (pH 5.5) for 30 min at 37°C. To form nucleosides, equimolar amount of 50 mM ammonium acetate (pH 9.3) and 2 units of Shrimp alkaline phosphatase (SAP, New England BioLabs, #M0371L) were added. The mixture was incubated for another 30 min at 37°C. Afterwards, the digested RNA was filtered through a 10 kDa Vivacon 500 filter (Sartorius, #VN01H02) at 14 000× g for 15 min, transferred to a plastic HPLC vial and analysed by LC-MS. For samples used in calibration of SELECT method, 1 ng of IVT-CEVd sample was mixed with 0.1 U of NuP1. In the next step 0.002 U of rSAP and and mixture of isotopically labeled standards Adenosine-13C5 (13C5A, #TRC-A280402, LGC) and *N6*-Methyladenosine-d3 (D3m^6^A, #TRC-M275897, LGC) with final 500 pM conentration was used. The rest of the protocol is the same as above.

**RNase A digestion**

200-300 ng of cCEVd or IVT-CEVd (70% m^6^A) were mixed with 100 ng of RNase A (Thermo Scientific, #EN0531) in 20 mM ammonium acetate buffer (pH 7.5) and incubated for 1h at 37°C. Volume of the mixture was 40 µL. Right after, two equivalents of acetonitrile were added and the sample was transferred to a plastic HPLC vial and analysed by LC-MS.

**RNase T1 digestion**

Firstly, the RNA (200-300 ng of cCEVd or IVT-CEVd 70% m^6^A) was denaturated in 50 µL of reaction containing 4M urea (Sigma-aldrich, #U5378), 20 mM ammonium acetate buffer (pH 7.5) and 100 µM EDTA (Sigma-aldrich, #E5134) by incubating for 5 min at 75°C. Right after, the mixture was placed on ice for
1 min and purified with a 10 kDa Vivacon 500 filter unit at 7500 ×g for 10 min at 4°C. To remove all remaining urea, one extra wash with 50 µL of molecular biology grade water (Sigma-aldrich, #95284) was performed. Filter was turned around and RNA was collected to new vial by centrifugation at 1000 ×g for 2 min at 4°C. The collected RNA was digested with 1000 units of RNase T1 (Thermo Scientific, #EN0542) in 20 mM ammonium acetate buffer (pH 7.5) containing 100 µM EDTA. The reaction (40 µL) was incubated for 1h at 37°C, mixed with two equivalents of acetonitrile and transferred to a plastic HPLC vial for subsequent LC-MS analysis.

**Liquid chromatography–mass spectrometry (LC-MS) analysis**

For the analysis, an HPLC (Acquity H-class, Waters) equipped with Xbridge Premier BEH amide column (2.5 µm, 4.6 mm X 150 mm, Waters) and an Acquity Premier HSST3 (1.7 µm, 2.1 mm X 150 mm, Waters) column were used for RNase and NuP1 digested RNAs, respectively. Mobile phase A (MP A) contained 10 mM ammonium acetate (Fisher, #A11550) in 90% acetonitrile (Fisher, #A955212). Mobile phase B (MP B) contained 10 mM ammonium acetate in ultrapure water (18.2 MΩ.cm, Purelab Chorus system, Elga). Autosampler was kept at 22°C. 10 µL of NuP1 and 50 µL of RNase-digested RNA samples were injected. The gradient of separation for each type of sample is shown in the Tables S2-S4. Mass spectrometry detection was performed with Xevo G2-XS QTof mass spectrometer (Waters) equipped with an electrospray ionization source. For all three types of digested RNAs, parameters were optimized to achieve the best sensitivity (Table S5). Samples used for calibration of SELECT method were analyzed using HPLC (Acquity Premier, Waters) equipped with Acquity Premier HSST3 (1.7 µm, 2.1 mm X 150 mm, Waters) column. MP A was acetoinitrile and MP B was ultrapure water with 0.1% formic acid (Fisher, #A117-50). Gradient of separation is shown in Table S2. Analysis was performed with Xevo TQ Absolute (Waters) triple-quadrupole mass spectrometer equipped with ESI source (ionization parameters were similar to those used in QTof). Absolute quantitation of adenosine (A) and *N6*-methyladenosine (m^6^A) was done by single point calibration method calculated from AUC (area under the curve) of particular analyte and its isotopically labeled standard. MRM (multiple reaction monitoring) transitions were following: A (268→136), 13C5A (273→136) m^6^A (282→150) and D3m^6^A (285→153).

**Oxford Nanopore Technology (ONT) direct RNA sequencing**

For the ONT direct RNA sequencing, three different RNA preparations isolated from infected plants were used: either double-PAGE-purified circular CEVd RNA (cCEVd #1 and #2) or CF11-RNAs enriched in highly structured RNAs digested with RNase R to remove linear RNAs (HS-cRNA #1). In addition, three in vitro synthetized circular RNAs were prepared as described above (IVT-CEVd #1-#3). The circular RNAs were linearized using mild ZnCl_2_-mediated fragmentation optimized to open the circular RNA at a single random position. 300-500 ng of purified circular CEVd RNA was mixed on ice with the linearization buffer (final concentration 2 mM ZnCl_2_, 10 mM Tris.Cl pH 8.0) in a total volume of 80 µL. The reaction was split into 4 x 20 µL aliquots in PCR tubes and incubated for 90 seconds at 75°C. Immediately after the incubation, the reaction was stopped by the addition of 2.2 µL of 0.5 M EDTA pH 8.0 (final concentration 50 mM) into each aliquot and putting on ice. The RNA was purified by Zymo Clean & Concentrator 5 and eluted in 22 µL of nuclease-free water. The ends were repaired by incubating the eluted linear RNA with 1 U Fast Alkaline Phosphatase (ThermoFisher #EF0651) in a total volume of 25 µL, incubated at 37°C for 10 minutes and stopped by addition of EDTA pH 8.0 to final concentration 50 mM. Afterwards, the RNA was purified with Zymo Clean & Concentrator 5 and eluted in 22 µL of water. The RNA was polyadenylated at the 3’ end using 1 U *E. coli* polyA polymerase (NEB #M0276) in a mixture containing 1 mM ATP and the *E. coli* polyA polymerase buffer in a total volume of 30 µL. Afterwards, the RNA was purified with Zymo Clean & Concentrator 5 and eluted in 12 µL of water. 9 µL (~180 ng) of polydenylated RNA was used for the SQK-RNA002 or SQK-RNA004 library preparation kit according to ONT protocol and sequenced on MinION Mk1C (Oxford Nanopore Technology). The summary of sequencing results is provided in Table S7.

**Direct RNA sequencing data basecalling and mapping of the reads.**

The direct RNA sequencing data generated with the RNA002 chemistry on 9.4.1 flowcell were processed with the ONT Guppy basecaller software on the MinION device, using High-accuracy settings (configuration file rna_r9.4.1_70bps_hac.cfg). The reads were mapped using minimap2 (version 2.24-r1122^5^) to the reference CEVd dimeric sequence prepared as a concatemer of two monomeric CEVd sequences to allow alignment of all fragments generated by non-specific cleavage of the original circular. The resulting reads with quality score < 10 and all non-primary alignments were removed from further analyses using samtools^6^.

**Analysis of RNA content for contaminating sequences**

Quality of Nanopore sequencing runs and libraries was evaluated by using the pycoqc program^7^ (v2.5.0.3). Reads with Qscore higher than 8 (to retain more reads and get more representative blast results) were used for minimap2 (v2.24-r1122) mapping^5^ against the ONT spike-in (ENO2) sequences and CEVd genome (GenBank ID S67438.1). Nanopore reads not mapping either on spike-in or on CEVd were filtered out for further analysis. Such reads were first annotated by blastn^8^ (v2.9.0) against nt database to verify their possible sequence homology with other known sequences, setting as threshold e-value 10^-7^ and saving the output in tabular format. Blastn annotated sequences were then aggregated and counted by keyword text-mining, including the expressions “exocortis”, “saccharomyces”, “viroid”, “polyprobe”, “ribosomal” and “rRNA”. The sequences not containing these keywords were considered as other annotations and further manually checked for possible relevant annotations or keywords. Blastn unannotated sequences were clustered using cd-hit^9^ (v4.8.1) at 85% of sequence identity to cover the nanopore technology error rate (10-15%). Finally, clustered sequences from the HS-RNA without RNase R treatment were used for a blastn search against the IVT clustered sequences (Table S7).

**Prediction of m^6^A positions by DRUMMER**

The data were prepared according to the developer instructions^10^ (<https://github.com/DepledgeLab/DRUMMER>) and DRUMMER was run with two replicates of each group (the cCEVd #1/HS-cRNA #1 vs IVT-CEVd #1/2 control) in the multiple comparison exome mode, m^6^A detection and, visualizations for individual transcripts set on. The results were filtered as follows: Adjusted p-value of the max.G_test < 0.05; abs(odds_ratio) > 1.5 ,max_odds_padj < 0.05, minimal coverage of 300 reads at position and an AC/AU motif within the 11-bp surrounding the predicted site. G-score was used for scoring the modification probability.

**Prediction of m^6^A positions by Nanocompore**

The filtered reads were processed using nanopolish (<https://github.com/jts/nanopolish>)^11^ followed by m^6^A prediction according to Nanocompore documentation at (<https://nanocompore.rna.rocks/>) using IVT-CEVd #1/IVT-CEVd #2 as duplicates of non-modified control; and cCEVd #1 and HS-cRNA #1 as duplicates of modified sample. The results were filtered for positions with -log_10_(GMM_pvalue) >2 (corresponding to pvalue < 0.01), and abs (Logit_LOR) value > 0.3 to increase sensitivity as suggested by Mulroney et al, 2023^12^. -log_10_(GMM_pvalue) was used for scoring the modification probability.

**Prediction of m^6^A positions by xPore**

The analysis by xPore was performed according to the documentation available at (<https://xpore.readthedocs.io/en/latest/index.html>)^13^. The analysis was run in the –diffmod mode, prefiltering method t-test at threshold 0.1, with duplicate IVT-CEVd #1 and IVT-CEVd #2 RNA samples assigned as KO and the duplicate cCEVd #1 and HS-cRNA #1 viroid RNA samples assigned as WT. The output (*majority_direction_kmer_diffmod.table*) was postfiltered by selecting for the adjusted p-value of the differential modification rate (pval_KO_vs_WT) < 0.05 and the 5-mer containing AC or AU motif. For scoring, the differential modification rate parameter was used.

**Prediction of m^6^A positions by Dorado base caller using the ONT RNA004 chemistry**

The circular RNAs (cCEVd #2 and IVT-CEVd #3) prepared to obtain a polyadenylated RNA as above were processed using the ONT SQK-RNA004 protocol and sequenced on the FLO-MIN004RA flowcell. The runs were performed for 3-5 hours producing between 500.000 and 1.000.000 raw reads. For the analysis of RNA content, the raw data was basecalled using High-Accuracy Basecaller (models hac). For the detection of the m^6^A modification, the .pod5 formatted raw data was basecalled using Dorado (version 0.5.2+7969fab, https://github.com/nanoporetech/dorado) with the dorado super high accuracy basecaller with m^6^A detection option (models sup, m6A_DRACH) using the rna004_130bps_sup@v3.0.1 and, mapping to the reference sequence (Figure S6A). Only primary mapping reads with quality score >10 were used for further analyses. To analyze the m^6^A predicted positions, a bed file containing the fraction modified value was generated by modkit (https://github.com/nanoporetech/modkit, version 0.2.4). Edge filter was used to remove first and last 10 read nucleotides in each read (--edge-filter 10,10). Differential base modification was assessed by modkit (modkit dmr pair) and dmr score was used for scoring differential modification.

**Detection of m^6^A modification by SELECT method**

The SELECT method was performed according to the original protocol^32^ with the following modifications: The calibration curve was constructed by mixing 0% and 100% m^6^A-containing IVT synthetized cCEVd to obtain 100%, 50%, 25%, 12.5% and 0% modified IVT CEVd. The content of m^6^A in the calibration samples was confirmed by LC-MS **(Figure S6).** The SELECT reaction was performed with the same conditions as described in the original protocol**^32^**, with 5 ng of CEVd RNA (calibration sample or cCEVd sample) per well. After the SELECT reaction (nick-sealing and ligation), the reaction mixture was diluted 5x with RNAse/DNAse-free water and 2 µL used for qPCR using primers SELECT-qPCR Fwd and SELECT-qPCR Rev. 2 µL of the diluted SELECT reaction was reverse transcribed using LunaScript® RT SuperMix (NEB #M3010) in a 10 µL reaction, which was subsequently diluted 6x with RNAse/DNAse-free water and used for qPCR using primers CEVd-qPCR Fwd and CEVd-qPCR Rev. 2 µL of either diluted SELECT or reverse transcription reaction was used per well in a 10 µL reaction volume of the Luna^®^ Universal qPCR Master Mix (NEB #M3003) containing 100 nM forward and reverse qPCR primers. PCR (1 min 95°C initial denaturation, 45 cycles of 15 seconds 95°C denaturation followed by 30 s 60 °C annealing/extension steps) with subsequent melting curve analysis (60°C to 95°C, ramping rate 0.11 °C /s) were performed in technical duplicate at the LightCycler® 480 Instrument II (Roche) and the data were analyzed using LightCycler® 480 software release 1.5.1.62. After the melting curve quality check, the Cq values were determined using the Abs quant/Fit points mode, the SELECT product Cq values were normalized by the CEVd RT-qPCR Cq values. The resulting ΔCq values of the calibration samples were plotted with the % m^6^A of the calibrator samples estimated by LC-MS (Figure S7 A), and the obtained linear regression was used to estimate the percentage m^6^A in the double-PAGE purified cCEVd samples measured at 1x and 4x dilution.

**Supplementary tables**

Table S 1: **Oligonucleotides used for full length CEVd cDNA amplification and cloning; oligonucleotides for SELECT method**

| **Name** | **Sequence (5' to 3')*^#^** |
| --- | --- |
| CEVd2.5phi-For1 | **TAATACGACTCACTATT**AGGTTCCTGTGGTGCTCACCTG |
| CEVd2.5phi-Rev2 | CAAGAAAGTTCCCGAGGGACC |
| CEVd2.5phi-For3 | **TAATACGACTCACTATT**AGGAGCTCGTCTCCTTCCTTT |
| CEVd2.5phi-Rev4 | GTTTCTCCGCTGGACGCCAGT |
| CEVd qPCR Fwd | CGCTGCTGGCTCCACATC |
| CEVd qPCR Rev | TTGGGGTTGAAGCTTCAGTTG |
| SELECT qPCR-Rev | ATGCAGCGACTCAGCCTCTG |
| SELECT qPCR-Fwd | TAGCCAGTACCGTAGTGCGTG |
| A6 SELECT 5’phos oligo | p-TCCCGAGGGACCCAGTCcagaggctgagtcgctgcat |
| A6 SELECT missing dT site oligo | tagccagtaccgtagtgcgtgCACAGGAACCTCAAGAAAG |
| A353 SELECT 5’phos oligo | p-TCCAAGGGCTTTCACCCTCcagaggctgagtcgctgcat |
| A353 SELECT missing dT site oligo | tagccagtaccgtagtgcgtgGAGGGACCCAGTCTAGGG |
| A360 SELECT 5’phos oligo | p-CTAGGGTTCCAAGGGCTTTCcagaggctgagtcgctgcat |
| A360 SELECT missing dT site oligo | tagccagtaccgtagtgcgtgAGTTCCCGAGGGACCCAG |
| * in bold is denoted the sequence of the T7 class II promoter phi2.5 | |

^#^ the lower case letters indicate the priming sites for SELECT qPCR primers

Table S 2: **HPLC gradient used in the analysis of NuP1 digested RNA.**

| **Time (min)** | **Flow (mL/min)** | **MP A (%)** | **MP B (%)** |
| --- | --- | --- | --- |
| 0 | 0.25 | 0 | 100 |
| 2 | 0.25 | 0 | 100 |
| 8 | 0.25 | 60 (15*) | 40 (80*) |
| 8.5 | 0.25 | 90 | 10 |
| 9.5 | 0.5 | 90 | 10 |
| 11 | 0.5 | 90 | 10 |
| 11.1 | 0.5 | 0 | 100 |
| 16 | 0.5 | 0 | 100 |
| 16.1 | 0.25 | 0 | 100 |
| 17 | 0.25 | 0 | 100 |

* Gradient for SELECT samples

Table S 3: **HPLC gradient used in the analysis of RNase A digested RNA.**

| **Time (min)** | **Flow (mL/min)** | **MP A (%)** | **MP B (%)** |
| --- | --- | --- | --- |
| 0 | 1 | 80 | 20 |
| 2 | 1 | 80 | 20 |
| 8 | 1 | 50 | 50 |
| 10 | 1 | 40 | 60 |
| 13 | 1 | 40 | 60 |
| 13.1 | 1 | 80 | 20 |
| 18 | 1 | 80 | 20 |
| 20 | 1 | 80 | 20 |

Table S 4: **HPLC gradient used in the analysis of RNase T1 digested RNA.**

| **Time (min)** | **Flow (mL/min)** | **MP A (%)** | **MP B (%)** |
| --- | --- | --- | --- |
| 0 | 1 | 70 | 30 |
| 2 | 1 | 70 | 30 |
| 16 | 1 | 40 | 60 |
| 20 | 1 | 40 | 60 |
| 20.1 | 1 | 70 | 30 |
| 25 | 1 | 70 | 30 |

Table S 5: **Electrospray ionization parameters used for the analysis of digested RNA.**

| **Parameters** | **NuP1** | **RNase A** | **RNase T1** |
| --- | --- | --- | --- |
| Ion mode | positive | positive | Negative |
| Mass range (m/z) | 50 - 600 | 300 - 1600 | 400 – 2000 |
| Capillary voltage (V) | 3000 | 3000 | 2200 |
| Sampling Cone (V) | 20 | 20 | 40 |
| Source Offset (V) | 40 | 40 | 40 |
| Source Temperature (°C) | 150 | 150 | 150 |
| Desolvation Temperature (°C) | 500 | 500 | 500 |
| Cone Gas (L/h) | 50 | 50 | 50 |
| Desolvation Gas (L/h) | 500 | 1000 | 1000 |
| Collision Energy | 4 | 6 | 6 |

Table S 6: **List of analyzed modified nucleosides in the NuP1 -LC-MS approach.**

| **Modification** | **molecular formula** | **retention time** | **charge** | **calculated m/z** | **measured m/z** | **mass error (ppm)** |
| --- | --- | --- | --- | --- | --- | --- |
| m^1^A | C11H15N5O4 | 2.32 | 1+ | 282.120 | 282.120 | 0.0 |
| A_m_ | C11H15N5O4 | 5.85 | 1+ | 282.120 | 282.121 | 3.5 |
| m^6^A | C11H15N5O4 | 6.15 | 1+ | 282.120 | 282.120 | 0.0 |
| m^6^A_m_ | C12H17N5O4 | 6.36 | 1+ | 296.136 | 296.136 | 0.0 |
| m^3^C | C10H15N3O5 | 2.02 | 1+ | 258.109 | 258.109 | 0.0 |
| m^5^C | C10H15N3O5 | 2.69 | 1+ | 258.109 | 258.108 | -3.9 |
| C_m_ | C10H15N3O5 | 2.71 | 1+ | 258.109 | 258.109 | 0.0 |
| m^2^G | C11H15N5O5 | 4.61 | 1+ | 298.117 | 298.115 | -6.7 |
| G_m_ | C11H15N5O5 | 4.74 | 1+ | 298.117 | 298.113 | -13.4 |
| m^7^G | C11H15N5O5 | 5.07 | 1+ | 298.117 | 298.114 | -10.1 |
| I | C10H12N4O5 | 2.71 | 1+ | 269.089 | 269.087 | -7.4 |

Table S 7: **Summary of ONT direct sequencing runs and their mapping.**

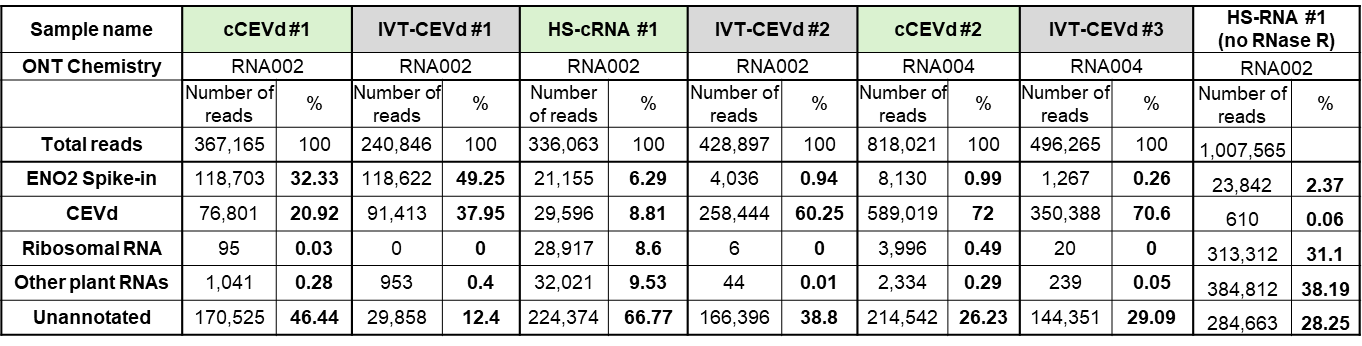

Table S 8: **List of analyzed oligonucleotides in the RNase T1 LC-MS experiment.**

| **Position**  **of A** | **Fragment** | **molecular formula** | **retention time** | **charge** | **calculated m/z** | **measured m/z** | **Mass error (ppm)** |
| --- | --- | --- | --- | --- | --- | --- | --- |
| **6** | AACUUUCUUG | C93H117N31O74P10 | 11.85 | 3- | 1052.783 | 1052.785 | 2.0 |
| **32** | CUCACCUG | C74H96N26O58P8 | 11.08 | 3- | 840.434 | 840.435 | 1.2 |
| **37** | ACCCUG | C56H73N21O43P6 | 9.66 | 3- | 636.745 | 636.747 | 3.1 |
| **90** | AUCCCCG | C65H85N24O50P7 | 10.34 | 3- | 738.426 | 738.425 | -1.4 |
| **102** | AAACCUG | C67H85N28O48P7 | 9.75 | 3- | 754.433 | 754.432 | -1.3 |
| **129** | ACAG | C39H50N18O27P4 | 7.10 | 2- | 662.094 | 662.094 | 0.0 |
| **151** | AUCACUG | C66H84N25O50P7 | 9.94 | 3- | 746.758 | 746.754 | -5.4 |
| **154** | AUCACUG | C66H84N25O50P7 | 9.94 | 3- | 746.758 | 746.754 | -5.4 |
| **173** | AAACAG | C59H74N28O39P6 | 8.43 | 3- | 660.429 | 660.430 | 1.5 |
| **209** | CUCCACAUCCG | C102H132N37O78P11 | 12.29 | 3- | 1153.479 | 1153.482 | 2.6 |
| **211** | CUCCACAUCCG | C102H132N37O78P11 | 12.29 | 3- | 1153.479 | 1153.482 | 2.6 |
| **216** | AUCG | C38H49N15O29P4 | 7.55 | 2- | 650.580 | 650.577 | -4.6 |
| **264** | ACUACCCG | C75H97N29O56P8 | 10.71 | 3- | 848.110 | 848.108 | -2.4 |
| **267** | ACUACCCG | C75H97N29O56P8 | 10.71 | 3- | 848.110 | 848.108 | -2.4 |
| **278** | AAACAACUG | C87H109N38O60P9 | 10.52 | 3- | 973.802 | 973.804 | 2.1 |
| **281** | AAACAACUG | C87H109N38O60P9 | 10.52 | 3- | 973.802 | 973.804 | 2.1 |
| **293** | CUUCAACCCCAAG | C122H156N47O90P13 | 12.70 | 4- | 1029.384 | 1029.378 | -5.8 |
| **312** | CUUUUCUUAUAUCUUCACUG | C184H231N57O149P20 | ND | 4- | 1559.417 | ND | N/A |
| **314** | CUUUUCUUAUAUCUUCACUG | C184H231N57O149P20 | ND | 4- | 1559.417 | ND | N/A |
| **320** | CUUUUCUUAUAUCUUCACUG | C184H231N57O149P20 | ND | 4- | 1559.417 | ND | N/A |
| **353** | AACCCUAG | C76H97N31O55P8 | 10.37 | 3- | 856.114 | 856.117 | 3.5 |
| **360** | ACUG | C38H49N15O29P4 | 7.55 | 2- | 650.580 | 650.577 | -4.6 |

| **Position of A** | **Fragment** | **molecular formula** | **retention time** | **charge** | **calculated m/z** | **measured m/z** | **mass error (ppm)** |
| --- | --- | --- | --- | --- | --- | --- | --- |
| **6** | Am6ACUUUCUUG | C94H119N31O74P10 | ND | 3- | 1057.455 | ND | N/A |
| **32** | CUCm6ACCUG | C75H98N26O58P8 | 10.97 | 3- | 845.106 | 845.106 | 0.0 |
| **37** | m6ACCCUG | C57H75N21O43P6 | 9.45 | 3- | 641.417 | 641.415 | -3.1 |
| **90** | m6AUCCCCG | C66H87N24O50P7 | ND | 3- | 743.098 | ND | N/A |
| **102** | AAm6ACCUG | C68H87N28O48P7 | 9.47 | 3- | 759.105 | 759.106 | 1.3 |
| **129** | m6ACAG | C40H52N18O27P4 | 6.64 | 2- | 669.102 | 669.103 | 1.5 |
| **151** | m6AUCACUG | C67H86N25O50P7 | 9.62 | 3- | 751.430 | 751.427 | -4.0 |
| **154** | AUCm6ACUG | C67H86N25O50P7 | 9.62 | 3- | 751.430 | 751.427 | -4.0 |
| **173** | AAm6ACAG | C60H76N28O39P6 | 8.10 | 3- | 665.100 | 665.100 | 0.0 |
| **209** | CUCCm6ACAUCCG | C103H134N37O78P11 | ND | 3- | 1158.151 | ND | N/A |
| **211** | CUCCACm6AUCCG | C103H134N37O78P11 | ND | 3- | 1158.151 | ND | N/A |
| **216** | m6AUCG | C39H51N15O29P4 | 7.19 | 2- | 657.588 | 657.589 | 1.5 |
| **264** | m6ACUACCCG | C76H99N29O56P8 | ND | 3- | 852.782 | ND | N/A |
| **267** | ACUm6ACCCG | C76H99N29O56P8 | ND | 3- | 852.782 | ND | N/A |
| **278** | AAm6ACAACUG | C88H111N38O60P9 | 10.31 | 3- | 978.474 | 978.479 | 5.1 |
| **281** | AAACAm6ACUG | C88H111N38O60P9 | 10.31 | 3- | 978.474 | 978.479 | 5.1 |
| **293** | CUUCAm6ACCCCAAG | C123H158N47O90P13 | ND | 4- | 1032.888 | ND | N/A |
| **314** | CUUUUCUUm6AUAUCUUCACUG | C186H233N57O149P20 | ND | 4- | 1565.921 | ND | N/A |
| **314** | CUUUUCUUAUm6AUCUUCACUG | C186H233N57O149P20 | ND | 4- | 1565.921 | ND | N/A |
| **320** | CUUUUCUUAUAUCUUCm6ACUG | C186H233N57O149P20 | ND | 4- | 1565.921 | ND | N/A |
| **353** | Am6ACCCUAG | C77H99N31O55P8 | ND | 3- | 860.786 | ND | N/A |
| **360** | m6ACUG | C39H51N15O29P4 | 7.19 | 2- | 657.588 | 657.589 | 1.5 |

Table S 9: **List of analyzed oligonucleotides in the RNase A LC-MS experiment.**

| **Position of A** | **Fragment** | **molecular formula** | **retention time** | **charge** | **calculated m/z** | **measured m/z** | **mass error (ppm)** |
| --- | --- | --- | --- | --- | --- | --- | --- |
| **6** | GGGAAC | C59H74N28O41P6 | 8.96 | 2+ | 1009.157 | 1009.160 | 3.0 |
| **32** | AC | C19H26N8O14P2 | 7.02 | 1+ | 653.112 | 653.113 | 1.5 |
| **37** | GAC | C29H38N13O21P3 | 7.82 | 1+ | 998.160 | 998.151 | -9.0 |
| **90** | AGGGAU | C59H73N27O42P6 | 8.93 | 2+ | 1009.649 | 1009.654 | 5.0 |
| **102** | GGGGAAAC | C79H98N38O54P8 | 9.32 | 2+ | 1346.207 | 1346.172 | -26.0 |
| **129** | GGGGGGGAC | C89H110N43O63P9 | 9.84 | 2+ | 1534.720 | 1534.678 | -27.4 |
| **151** | GGAU | C39H49N17O29P4 | 8.40 | 1+ | 1344.191 | 1344.175 | -11.9 |
| **154** | AC | C19H26N8O14P2 | 7.02 | 1+ | 653.112 | 653.113 | 1.5 |
| **173** | GGAGAAAC | C79H98N38O53P8 | 9.14 | 2+ | 1338.210 | 1338.201 | -6.7 |
| **209** | AC | C19H26N8O14P2 | 7.02 | 1+ | 653.112 | 653.113 | 1.5 |
| **211** | AU | C19H25N7O15P2 | 6.80 | 1+ | 654.096 | 654.096 | 0.0 |
| **216** | GAU | C29H37N12O22P3 | 7.74 | 1+ | 999.140 | 999.144 | 4.0 |
| **264** | GGAGAC | C59H74N28O41P6 | 8.96 | 2+ | 1009.157 | 1009.160 | 3.0 |
| **267** | AC | C19H26N8O14P2 | 7.02 | 1+ | 653.112 | 653.113 | 1.5 |
| **278** | GGAAAC | C59H74N28O40P6 | 8.76 | 2+ | 1001.160 | 1001.147 | -13.0 |
| **281** | AAC | C29H38N13O20P3 | 7.47 | 1+ | 982.165 | 982.161 | -4.1 |
| **293** | AAC | C29H38N13O20P3 | 7.47 | 1+ | 982.165 | 982.161 | -4.1 |
| **312** | AU | C19H25N7O15P2 | 6.80 | 1+ | 654.096 | 654.096 | 0.0 |
| **314** | AU | C19H25N7O15P2 | 6.80 | 1+ | 654.096 | 654.096 | 0.0 |
| **320** | AC | C19H26N8O14P2 | 7.02 | 1+ | 653.112 | 653.113 | 1.5 |
| **353** | GGAAC | C49H62N23O34P5 | 8.61 | 2+ | 836.634 | 836.632 | -2.4 |
| **360** | AGAC | C39H50N18O27P4 | 8.10 | 2+ | 664.110 | 664.110 | 0.0 |

| **Position of A** | **Fragment** | **molecular formula** | **retention time** | **charge** | **calculated m/z** | **meassured m/z** | **mass error (ppm)** |
| --- | --- | --- | --- | --- | --- | --- | --- |
| **5** | GGGAm6AC | C60H76N28O41P6 | 8.86 | 2+ | 1016.165 | 1016.160 | -4.9 |
| **32** | m6AC | C20H28N8O14P2 | 6.73 | 1+ | 667.128 | 667.130 | 3.0 |
| **37** | Gm6AC | C30H40N13O21P3 | 7.68 | 1+ | 1012.175 | 1012.172 | -3.0 |
| **90** | AGGGm6AU | C60H75N27O42P6 | 8.77 | 2+ | 1016.657 | 1016.656 | -1.0 |
| **102** | GGGGAAm6AC | C80H100N38O54P8 | 9.24 | 2+ | 1353.215 | 1353.194 | -15.5 |
| **129** | GGGGGGGm6AC | C90H112N43O63P9 | 9.74 | 2+ | 1541.733 | 1541.692 | -26.6 |
| **151** | GGm6AU | C40H51N17O29P4 | 8.23 | 1+ | 1358.207 | 1358.197 | -7.4 |
| **154** | m6AC | C20H28N8O14P2 | 6.73 | 1+ | 667.128 | 667.130 | 3.0 |
| **173** | GGAGAAm6AC | C80H100N38O53P8 | 9.07 | 2+ | 1345.218 | 1345.203 | -11.2 |
| **209** | m6AC | C20H28N8O14P2 | 6.73 | 1+ | 667.128 | 667.130 | 3.0 |
| **211** | m6AU | C20H27N7O15P2 | 6.49 | 1+ | 668.112 | 668.109 | -4.5 |
| **216** | Gm6AU | C30H39N12O22P3 | 7.59 | 1+ | 1013.159 | 1013.162 | 3.0 |
| **264** | GGAGm6AC | C60H76N28O41P6 | 8.86 | 2+ | 1016.165 | 1016.160 | -4.9 |
| **267** | m6AC | C20H28N8O14P2 | 6.73 | 1+ | 667.128 | 667.130 | 3.0 |
| **278** | GGAAm6AC | C60H76N28O40P6 | 8.67 | 2+ | 1008.168 | 1008.165 | -3.0 |
| **281** | Am6AC | C30H40N13O20P3 | 7.29 | 1+ | 996.180 | 996.174 | -6.0 |
| **293** | Am6AC | C30H40N13O20P3 | 7.29 | 1+ | 996.180 | 996.174 | -6.0 |
| **211** | m6AU | C20H27N7O15P2 | 6.49 | 1+ | 668.112 | 668.109 | -4.5 |
| **211** | m6AU | C20H27N7O15P2 | 6.49 | 1+ | 668.112 | 668.109 | -4.5 |
| **320** | m6AC | C20H28N8O14P2 | 6.73 | 1+ | 667.128 | 667.130 | 3.00 |
| **353** | GGAm6AC | C50H64N23O34P5 | 8.61 | 2+ | 843.642 | 843.638 | -4.1 |
| **360** | m6AGAC | C40H52N18O27P4 | 7.96 | 2+ | 671.118 | 671.117 | -1.5 |

**Supplementary figures**

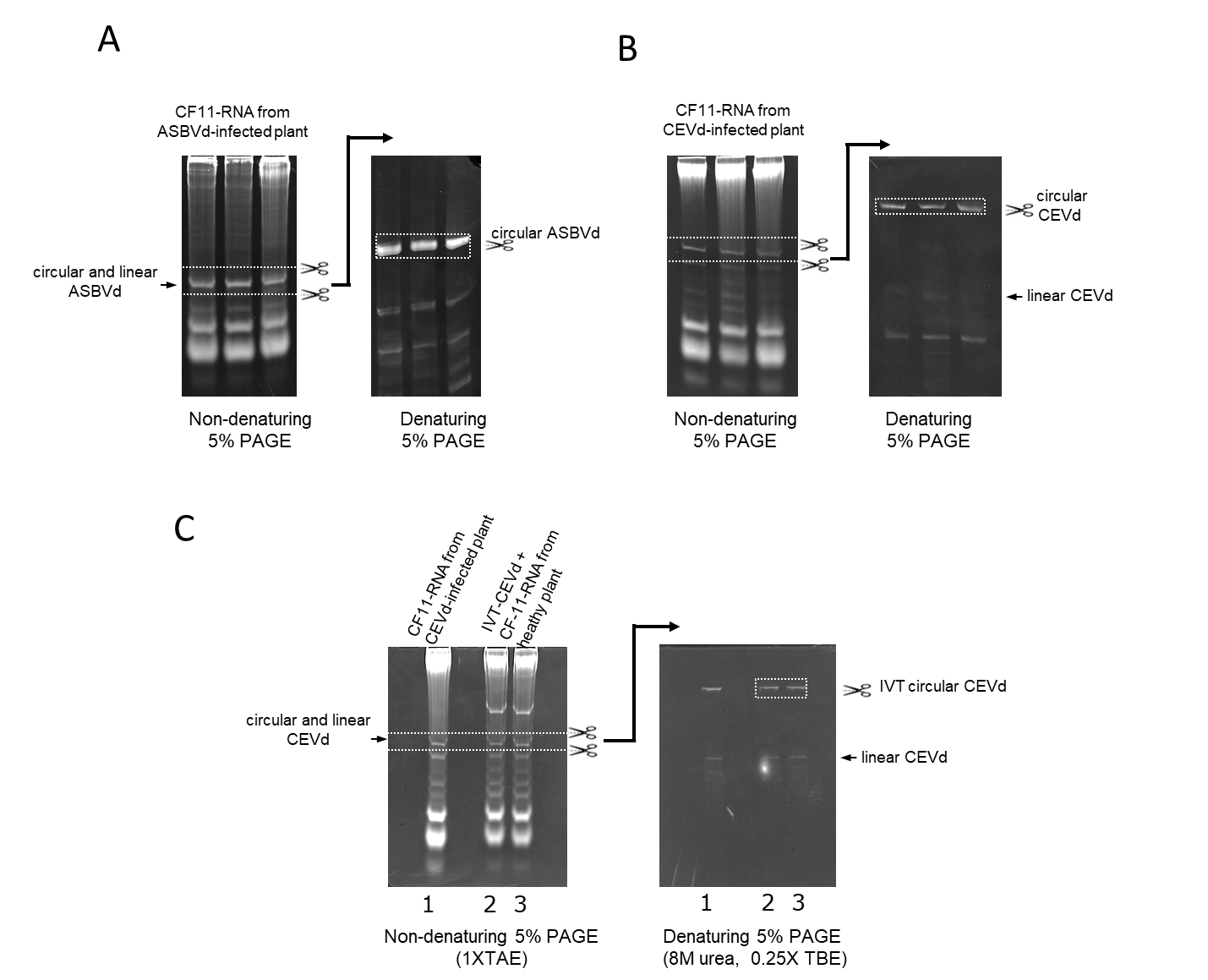

Figure S 1: **Purification of circular viroid RNA.** Double polyacrylamide gel electrophoresis (PAGE) system for purification of circular RNA forms of avocado sunblotch viroid (ASBVd) (panel A), citrus exocortis viroid (CEVd) (panel B) and in vitro synthesized circular CEVd RNA (IVT-CEVd). Nucleic acid preparations enriched in highly structured RNAs and obtained by CF11 chromatography in presence of 35% of ethanol (CF-11 RNAs) from ASBVd-infected avocado (panel A), CEVd-infected Gynura aurantiaca (panel B and line 1 panel C), and healthy G. aurantiaca spiked with IVT-CEVd (lines 2 and 3 of panel C) were fractionated in 5% PAGE under non-denaturing conditions (on the left of each panel). After ethidium bromide staining, the gel area within the dashed white lines was cut and applied on top of a denaturing 5% PAGE (on the right of each panel), which was subjected to electrophoresis and stained in the same way. Under denaturing conditions, the circular forms of the viroids migrate on the top of the gel, retarded with respect to the linear forms of the same size. The bands corresponding to the circular forms of the viroids that were cut from the gel and eluted are within a white-dashed square.

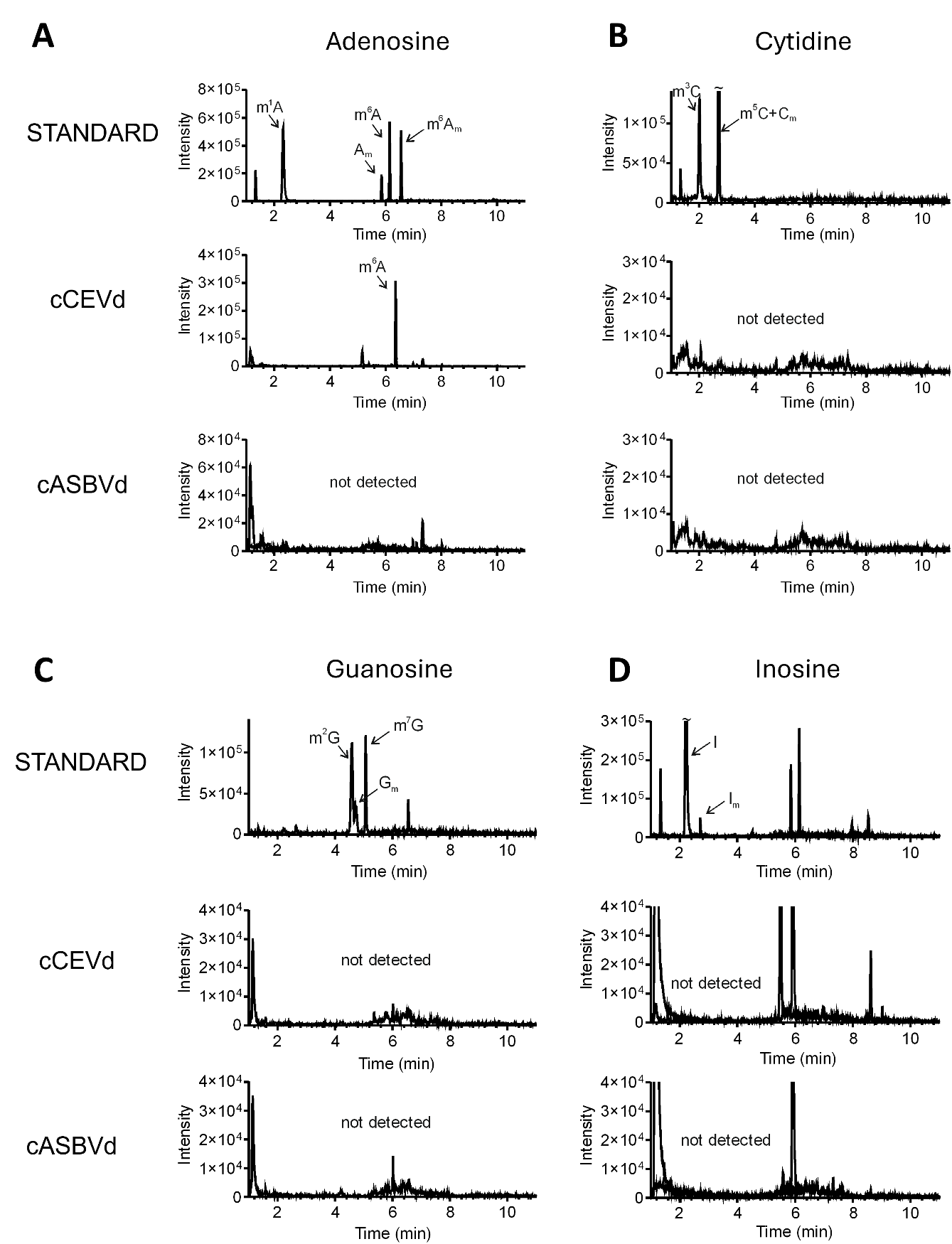

Figure S 2: **Extracted ion chromatograms of modified nucleosides in the NuP1/rSAP digested cCEVd and cASBVd.** (A) Combined EIC of m/z=282.120 and m/z=296.136 in standard mixture and viroid samples. (B) EIC of m/z=258.109 in standard mixture and viroid samples. (C) EIC of m/z=298.117 in standard mixture and viroid samples. (D) EIC of m/z=269.089 in standard mixture and viroid samples.

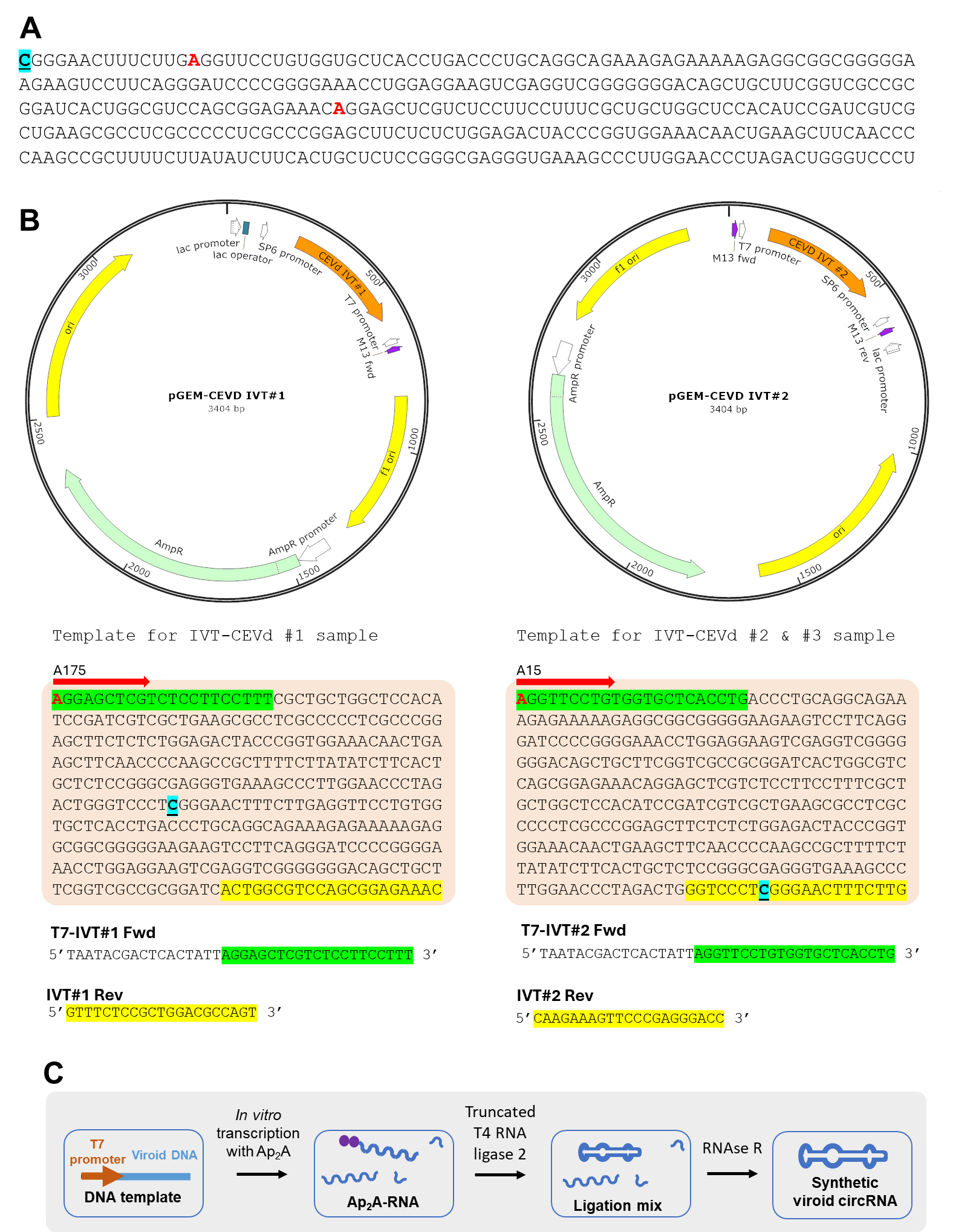

Figure S 3**: Plasmid templates for in vitro transcription (IVT).** (A) The CEVd sequence of G. aurantiaca isolate characterized in this work (GenBank ID PP446493). The blue-highlighted C marks the +1 position of the CEVd as reported in the CEVd RefSeq (ID NC_001464.1). The red-highlighted A letters indicate the start of transcription in the two different IVT templates. (B) The CEVd sequence was cloned into pGEM-T plasmid starting either from the A175 (plasmid IVT#1) or from the A15 (plasmid IVT#2) of the CEVd sequence (G. aurantiaca isolate). The orange-highlighted sequence is the sequence amplified from each plasmid. The red arrow indicates the start of T7 transcription, with the letter A175/A15 indicating the +1 transcribed base of the transcription relative to the reference sequence. Green highlighted sequence marks the priming site of the Fwd primer used for PCR amplification, the yellow highlighted sequence the priming site of the reverse primers. (C) A scheme of the preparation of circular IVT RNA samples.

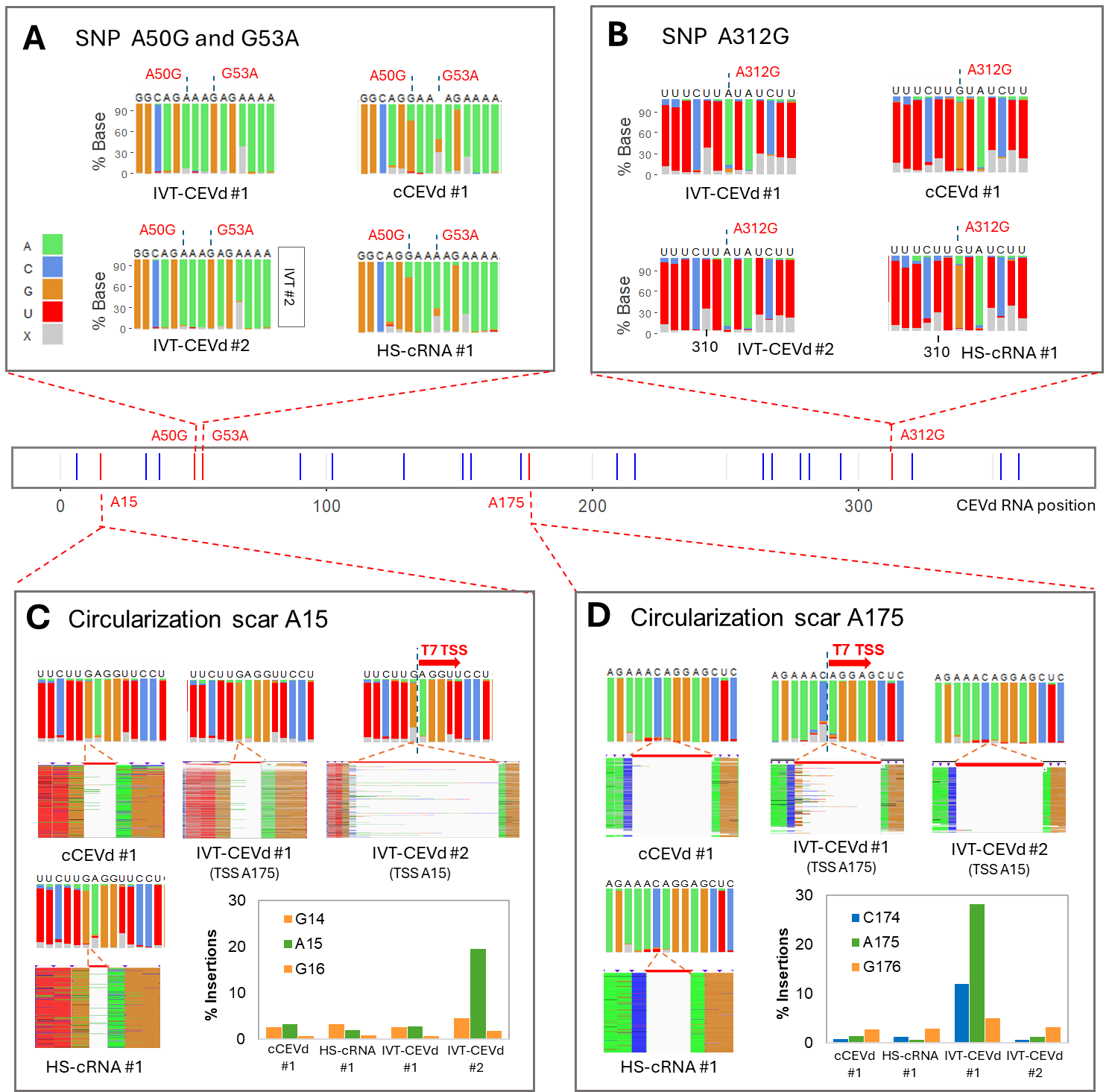

Figure S 4: **Artefacts in prediction of m^6^A.** (A) The single nucleotide polymorphisms A50G and G53A and (B) A312G were detected in the cCEVd #1/HS-cRNA #1 sample compared to the reference sequence. Due to differences in base frequencies at the polymorphic position, the nanopore signal is affected at this and surrounding positions, causing high modification scores in some algorithms. (C-D) The “circularization scars” A15 and A175 are artefacts occurring on the in vitro synthetized and circularized RNAs, due to the non-templated addition nucleotides by T7 polymerase at the 3’ end of the template. After circularization, these extra bases cause insertions in the mapped sequence reads, leading to a high score in some prediction algorithms. The changes in base-called nucleotides are visible at the ligation junction position 14/15 (reference numbering) in the IVT#2 transcribing from 5’ A15 and ending with 3’ G14 (C) This scar does not occur in the IVT#1 where the transcription starts at position A175. Conversely, the circularization scar in IVT#1 occurs at position C174/A175 and non-distorted signal is observed at A15 position. Expectedly, the A15 and A175 positions are unaffected in the cCEVd #1/HS-cRNA #1 samples.

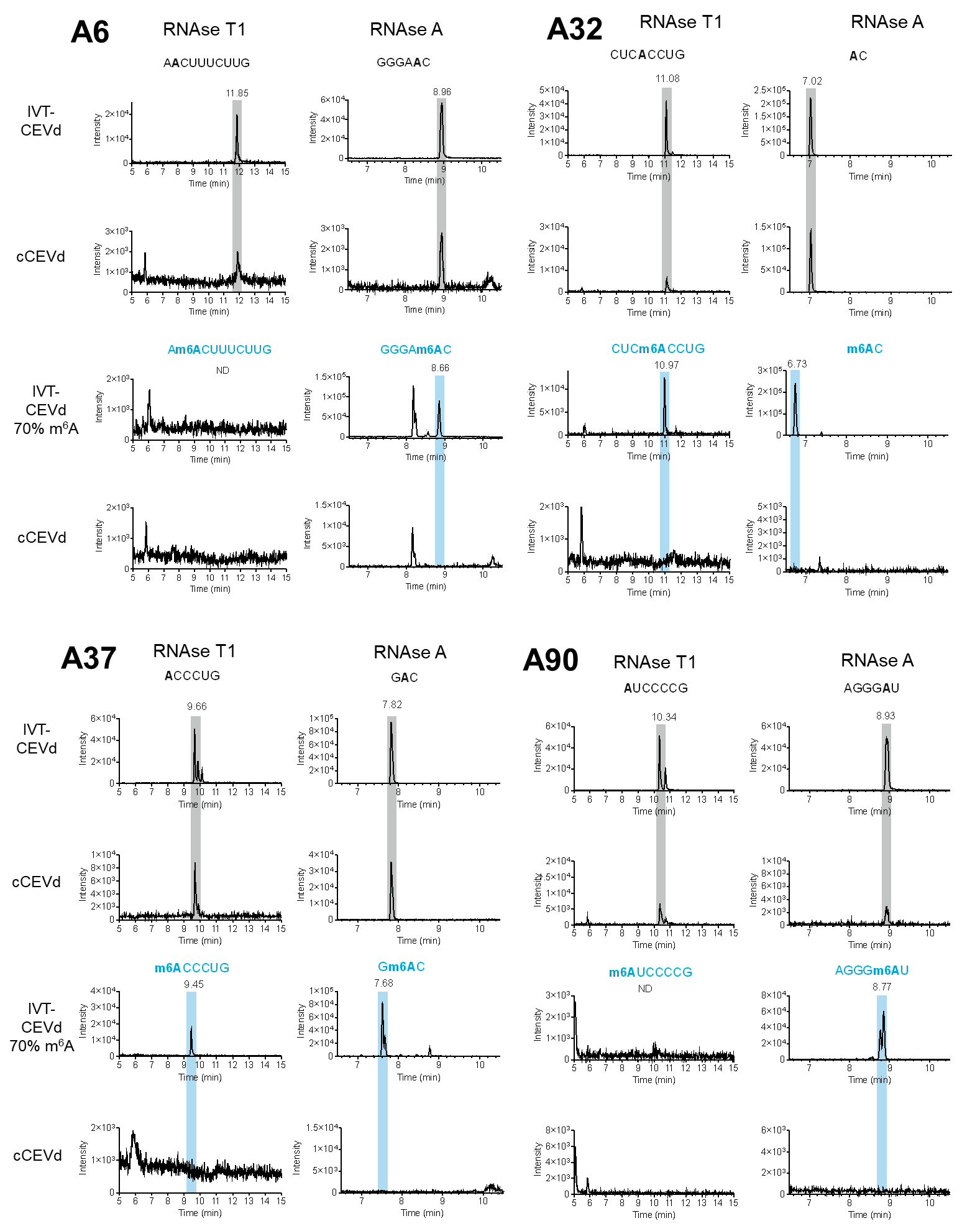

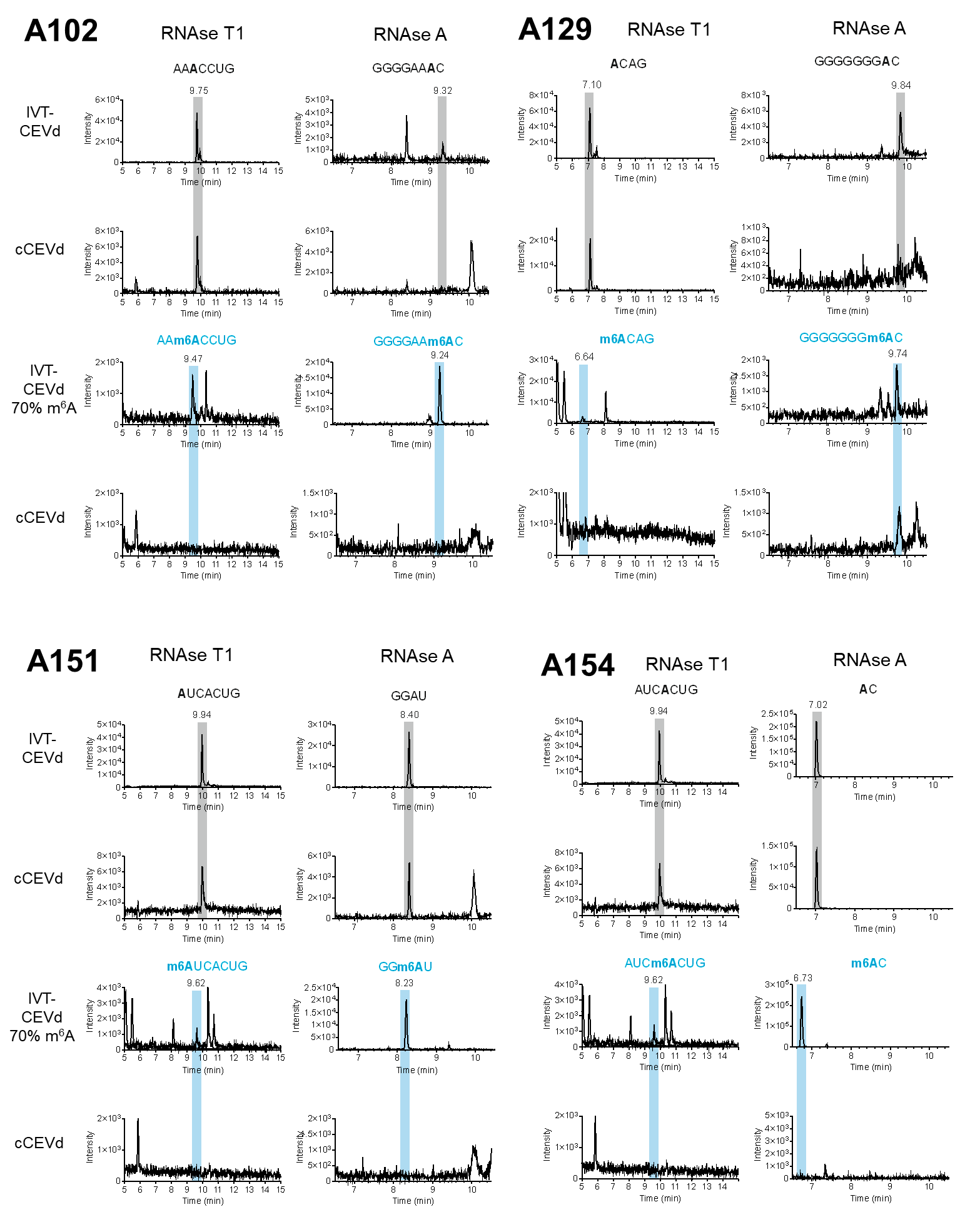

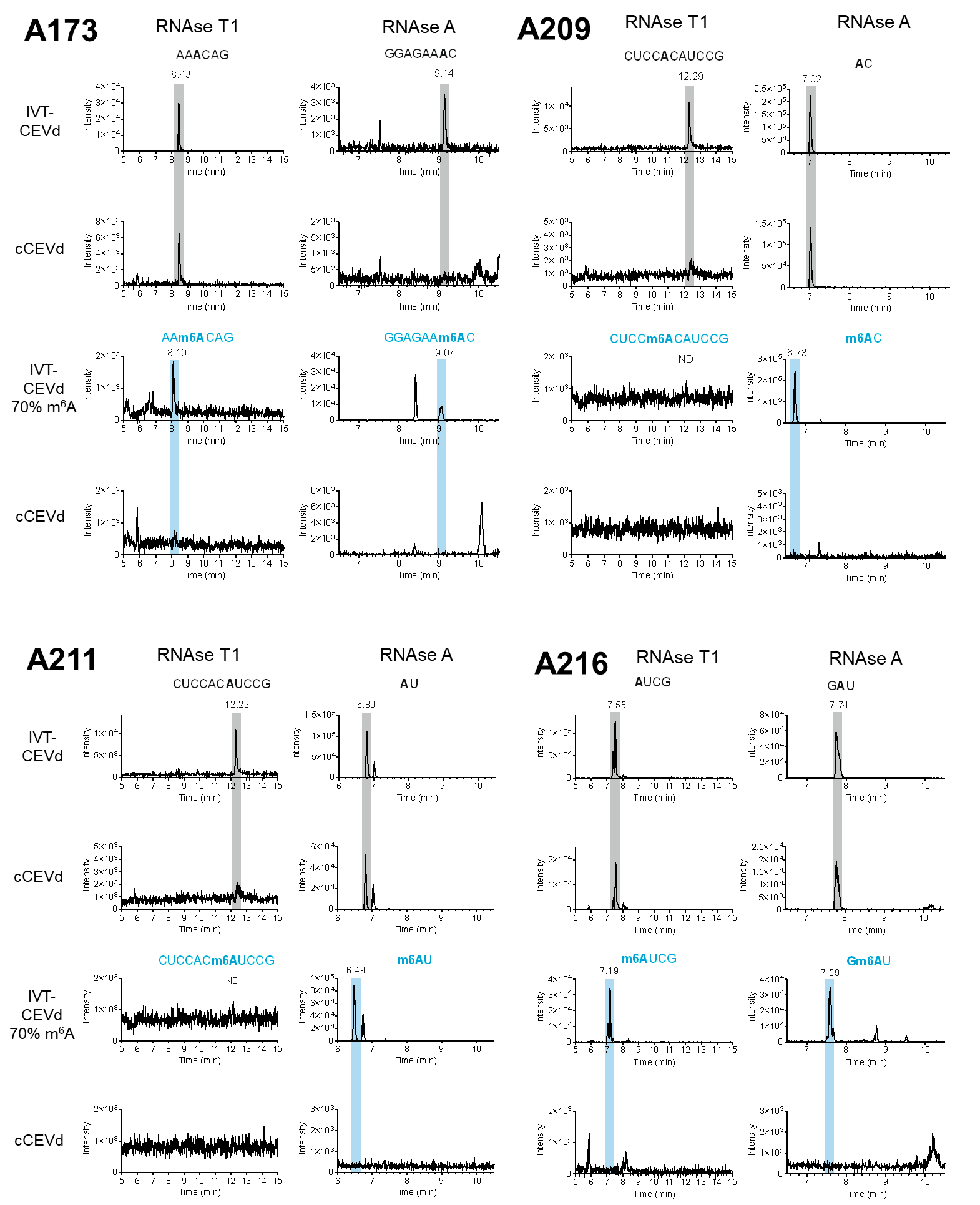

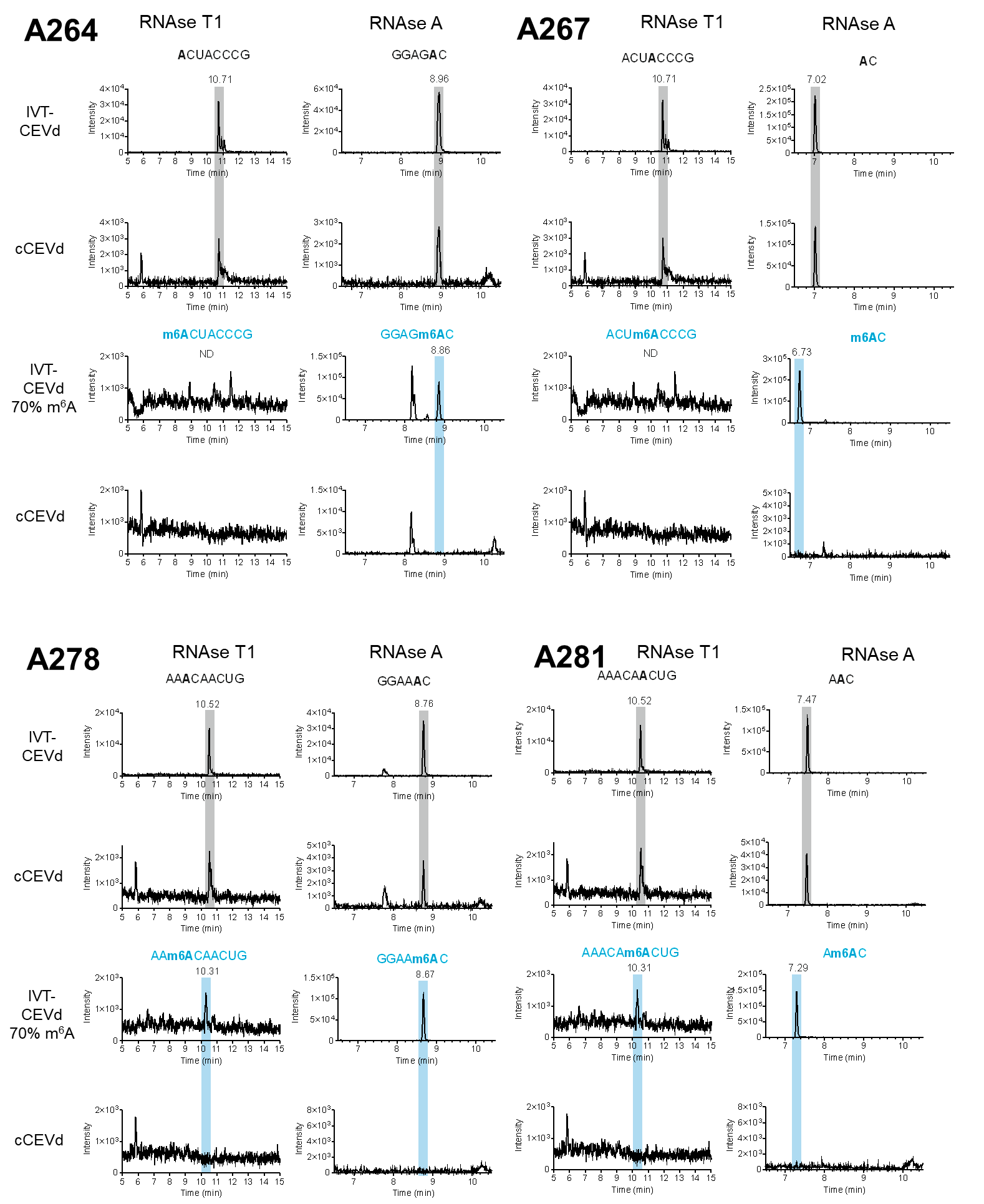

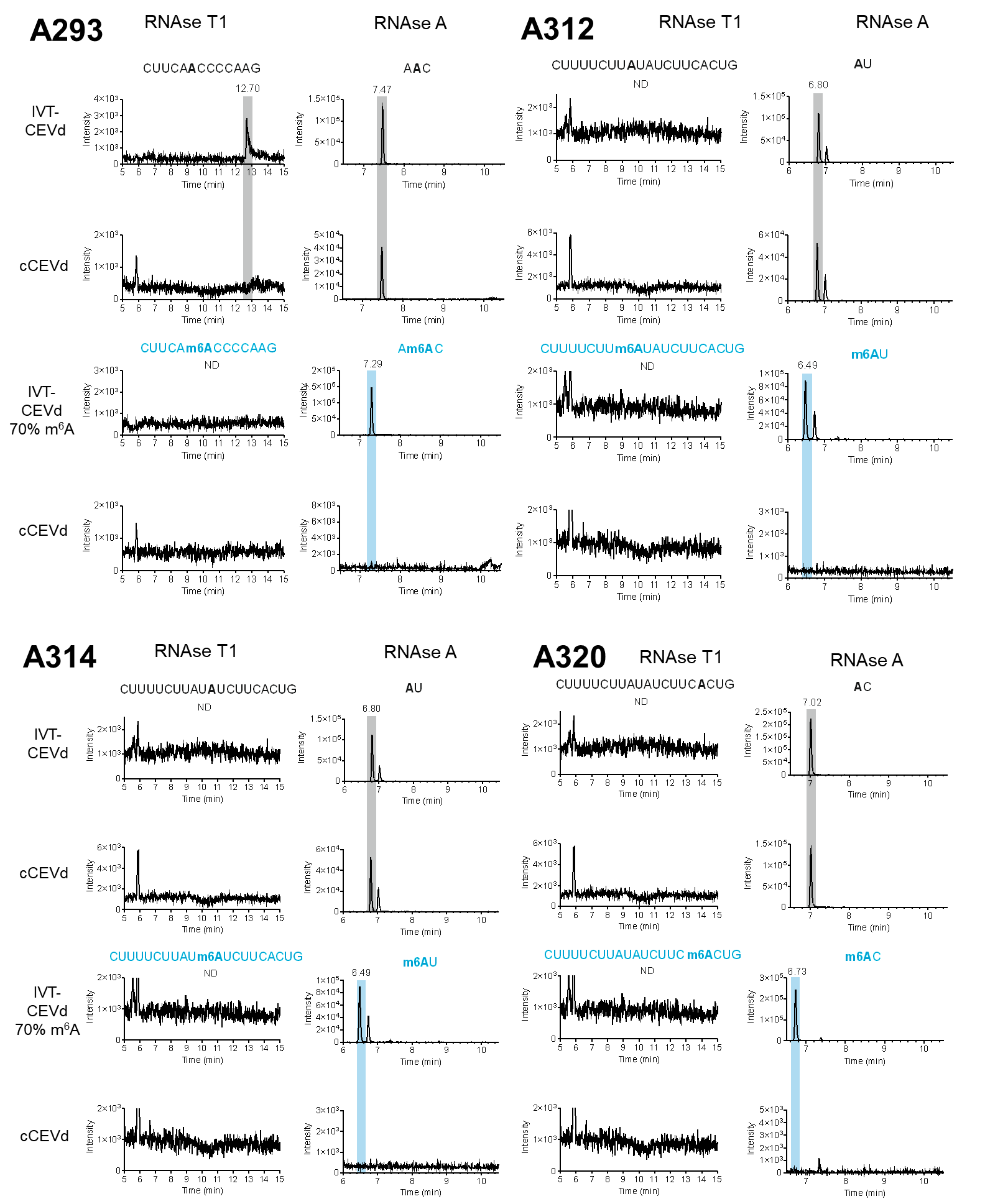

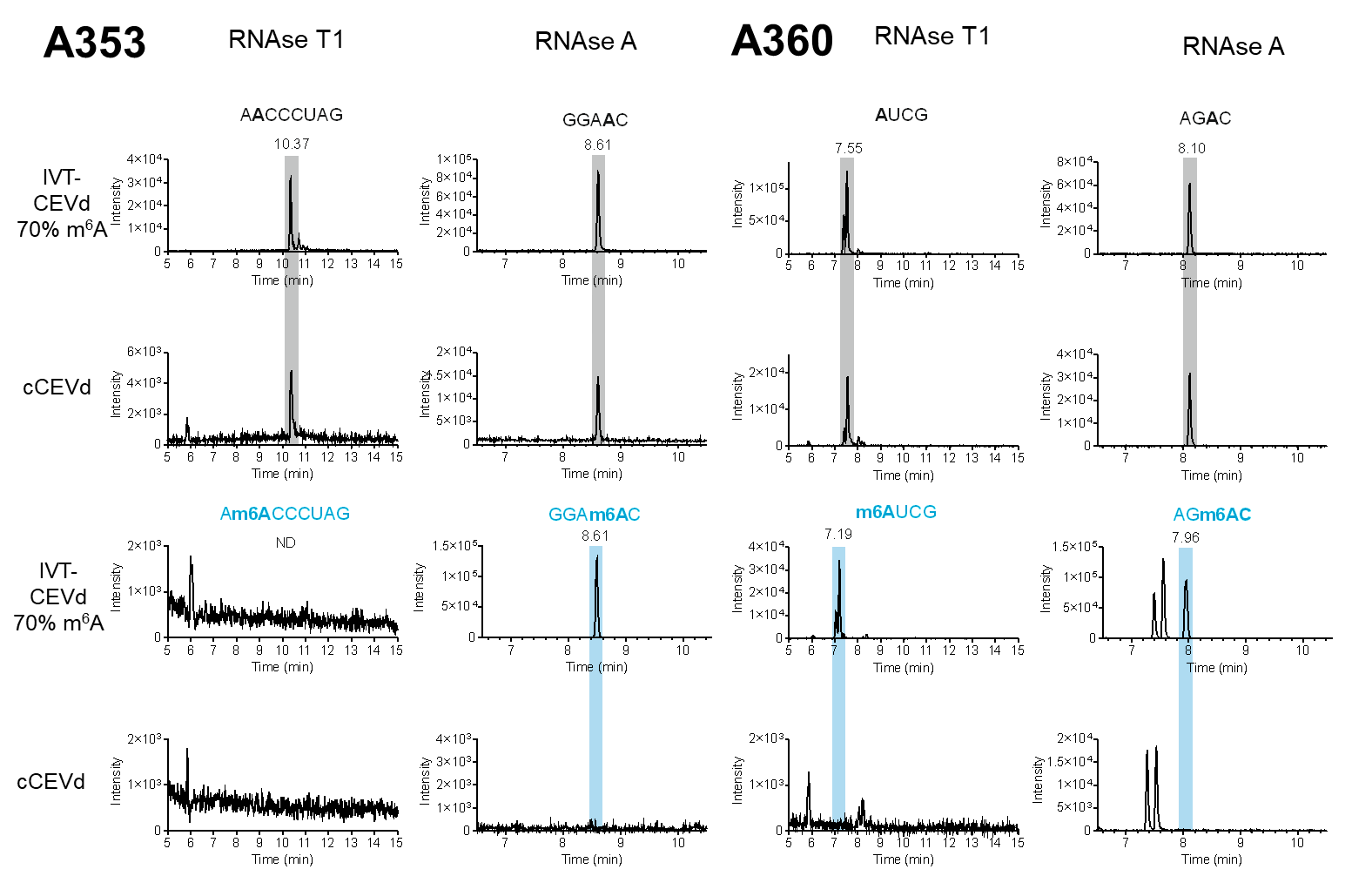

Figure S 5 (continued): Extracted ion chromatograms of analyzed oligonucleotides in RNase T1/RNase A LC-MS experiments for each AU/AC position within CEVd sequence in IVT CEVd 70% m^6^A and cCEVd samples. Grey rectangle marks retention time of unmethylated oligonucleotides and blue rectangle of methylated ones.

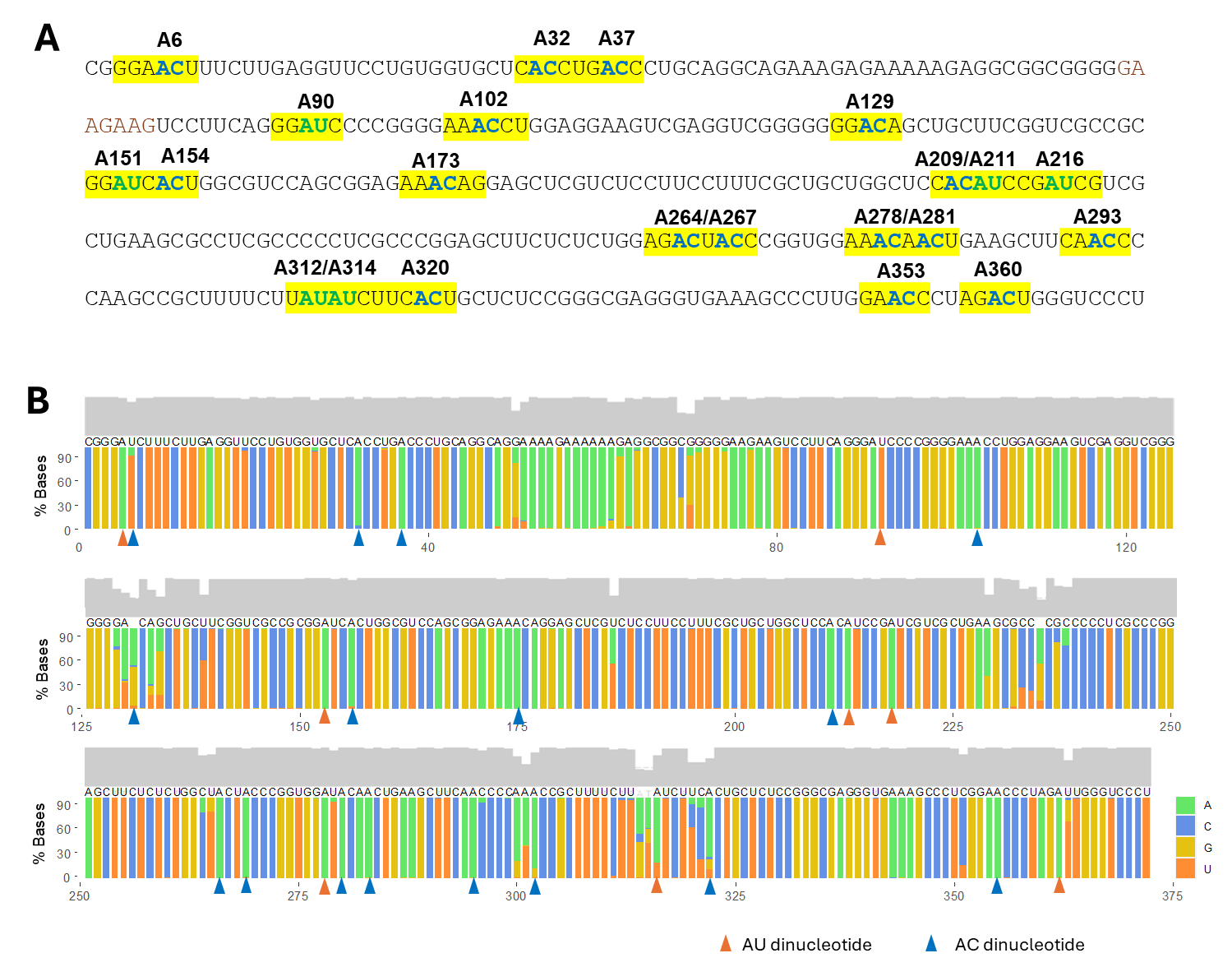

Figure S6: **Overview of AC/AU dinucleotides within CEVd sequence and their conservation in CEVd variants**. (A) The reference sequence with all AC/AU motifs that were analyzed using the RNAse T1/A – LC-MS approach. (B) The conservation of AC/AU dinucleotides in 148 CEVd sequences retrieved from public nucleotide databases (See Supplementary Excel file 1). Each nucleotide is color coded, and the bar shows the percentual proportions of each nucleotide at a given position based on multiple sequence alignment. The positions of AU (green circles) and AC (blue circles) are marked above the bar chart. The sequence above the barchart represents the consensus sequence. For positions where no nucleotide is present in >50% sequences, the nucleotide is left blank. The gray chart above the bar chart represents the conservation score.

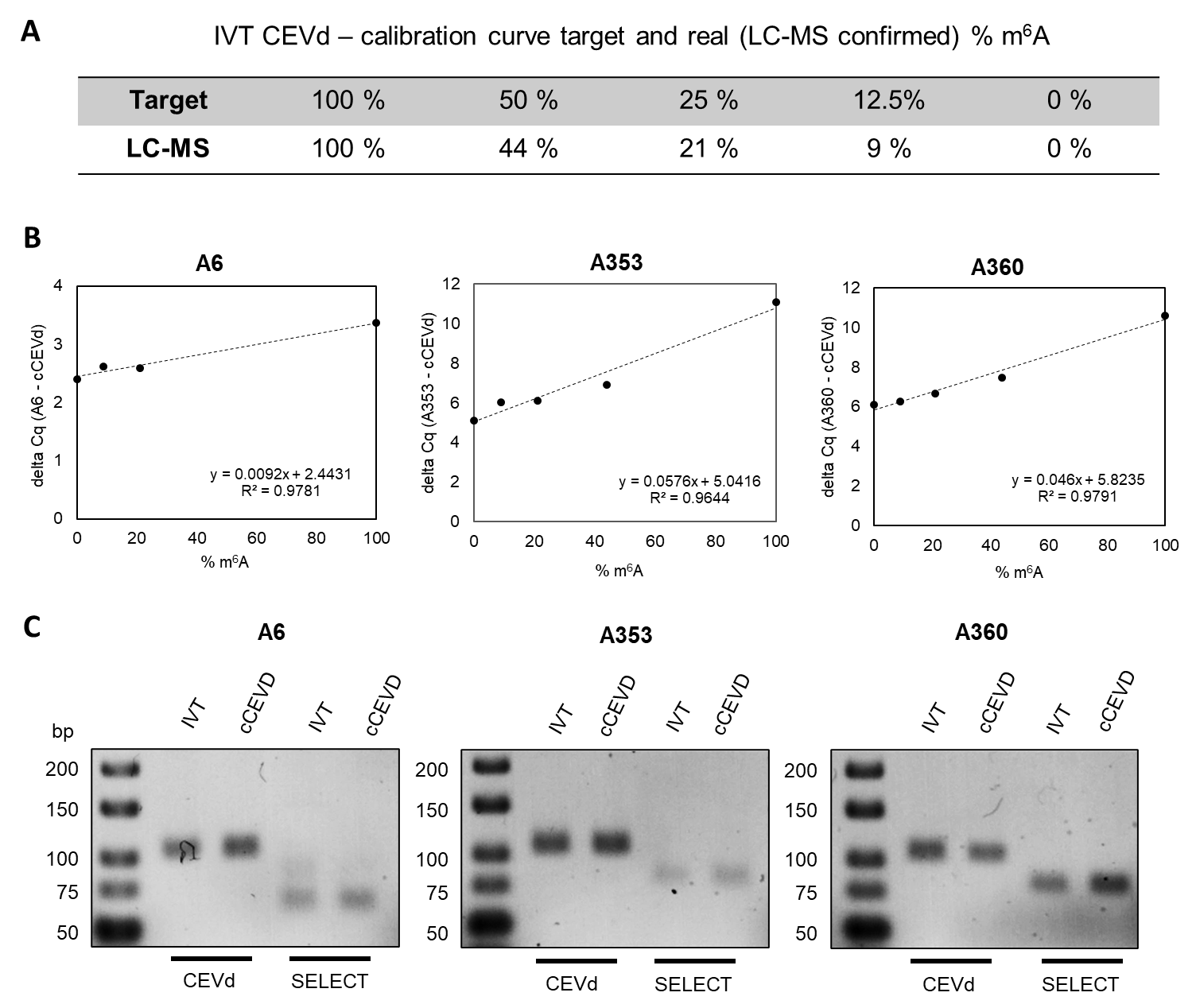

Figure S7: **Calibration curves for the SELECT method.** (A) The expected (upper row) versus measured (lower row) percentage in the calibration samples. For the calibration curve plotting and linear regression, the values measured by LC-MS were used. (B) Example calibration curves of the SELECT method for the investigated CEVd positions A6, A353 and A360.

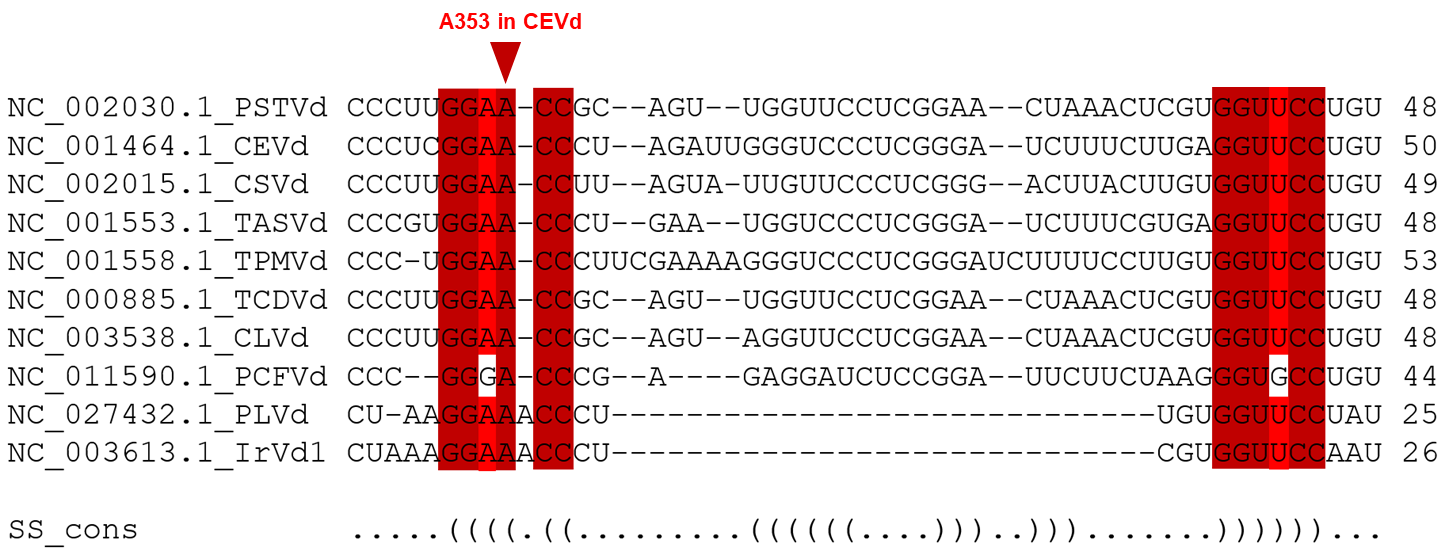

Figure S8: **Stockholm alignment of the left terminal region sequence of the members of the genus Pospiviroid.** For each species, the refseq variant has been reported together with its respective Genbank accession number. Red boxes indicate the conservation of the motifs in the alignment. Darker red indicates the highest conservation. At the bottom of the alignment, “SS cons” row shows the consensus secondary structure in WUSS notation.
